# Metagenomic Analyses Reveal a Constrained Network of Nutritional Symbionts in Kissing Bugs

**DOI:** 10.64898/2026.04.20.719636

**Authors:** G. Rignault, M. Merle, E. Folly-Ramos, C. E. Almeida, M. Harry, J. Filée

## Abstract

Triatominae bugs are the main vectors of Chagas disease in Latin America and rely on microbial nutritional symbiosis to complement their haematophagous diet with B-vitamins. While *Rhodococcus* bacteria have been identified as key symbionts, diverse metabarcoding analyses have suggested additional candidates. However, symbiont genomic data and metabolic capabilities remain largely uncharacterized. To address this gap, we generated metagenomic assemblies for 14 Triatominae and captured 15 bacterial genomes belonging to 4 genera (*Rhodococcus, Wolbachia, Symbiopectobacterium* and *Arsenophonus*) across 9 triatominae species. We identified five co-infection cases, including one involving two distinct *Arsenophonus* symbionts, one exhibiting hallmarks of massive genome degradation. Phylogenetic analyses revealed that Triatominae-associated symbionts form monophyletic groups within each genus, suggesting common origins followed by co-evolution with their hosts. Annotation of vitamin B metabolic genes indicates that most symbionts harbour incomplete pathways, with evidence of metabolic complementation between co-infecting symbionts. Additionally, we identified bacterial genes laterally transferred into host insect genomes, interpreted as footprints of present or past symbiotic associations. Nearly all Triatominae genomes displayed transferred genes from all four bacterial genera, including hosts with no detectable symbiont in genome assemblies. Taken together with these discoveries support the existence of a stable and limited network of four possible nutritional symbiont lineages with rare evidence of symbiont turn-overs.

**Significance statement:** Triatominae bugs, vectors of Chagas disease, are known to harbor a diverse community of nutritional bacterial symbionts whose genomic and metabolic roles have remained largely unexplored. By reconstructing 15 symbiont genomes that segregate as four bacterial genera, we provide important insight into the origins, the evolution and the metabolic structure of the nutritional symbiosis in triatominae. These findings support a stable, evolutionary conserved network of nutritional symbionts with limited turnover.

## Introduction

Triatominae bugs (Hemiptera: Reduviidae: Triatominae) are hemophagous insect vectors of *Trypanosoma cruzi*, the causative agent of Chagas disease which affects about 7 million people in Latin America (Ferreira et al., 2025). Also known as “kissing bugs”, triatominae compose a diversified group that have been divided into more than 15 genera and more than one hundred species (Schofield and Galvão, 2009) although species classification remains controversial (Filée et al., 2022).

Beyond their role as disease vectors, some triatominae species, such as *Rhodnius prolixus* have served as model insects for the study of hematophagy and host-symbiont interactions. The seminal work of Wigglesworth established that larval development of triatominae bugs critically depends on the presence of gut symbiotic bacteria (Wigglesworth, 1936). Further, Baines identified a midgut Actinobacteria, originally classified as *Nocardia rhodnii* and later reassigned to *Rhodococcus rhodnii*, and proposed that it supplies essential B-group vitamins, notably biotin, to its host (Baines, 1956). This interaction, known as “nutritional mutualism”, is widespread in insects and involves a wide range of microbial taxa and metabolic supplementation (Sudakaran et al., 2017).

The symbiont *Rhodococcus* is transmitted horizontally via egg-surface contamination or coprophagy (Wigglesworth, 1936). However, despite this long-standing paradigm, the precise metabolic benefits it provides to the host remain speculative (Gilliland et al., 2023). Numerous contradictory results suggest that the nutritional mutualism between *Rhodococcus* and *Rhodnius* is facultative rather than obligate, as its importance depends on rearing conditions, the host blood source, and the *R. rhodnii* strain involved. In some studies, larval development of symbiont-free insects is fully restored when blood meals are supplemented with B vitamins (Auden, 1974; Lake and Friend, 1968). In other studies, the requirement for such supplementation varies with the type of blood diet: *Rhodnius* individuals fed on mouse blood do not require B-vitamin supplementation, whereas supplementation is essential for bugs fed on rabbit blood (Baines, 1956; Nyirady, 1973). Moreover, host development was sometimes comparable using diverse *Rhodococcus* strains and/or auxotrophic mutants unable to produce B vitamins were used, suggesting that the symbiosis may involve additional metabolic functions beyond vitamin provisioning (Gilliland et al., 2023; Hill et al., 1976). In *Triatoma infestans*, aposymbiotic nymphs suffer a growth arrest even when their blood meal is supplemented with *R. rhodnii* but the development is restored when a *Corynebacterium* sp. is added to the food (Durvasula et al., 2008). These studies also raise the possibility that secondary symbionts may reinforce or compensate for the mutualistic relationships, as observed in other Hemiptera, including aphids (Manzano-Marín et al., 2023) and leafhoppers (Nishino et al., 2016). This suggests that triatominae symbioses may involve multi-partner interactions rather than a single obligate symbiont.

Recent advances in sequencing techniques have greatly expanded our knowledge of symbiont diversity in triatominae. In particular, microbial cultivation approaches and 16S rRNA-based metabarcoding studies have revealed a high diversity of gut symbionts shaped by multiple factors, including host species (Brown et al., 2018; Da Mota et al., 2012; Gumiel et al., 2015), geographic origin especially between wild and strain populations (Oliveira et al., 2018; Orantes et al., 2018; Rodríguez-Ruano et al., 2023), feeding status (Díaz et al., 2016; Gutiérrez-Millán et al., 2025; Rodríguez-Ruano et al., 2023), and *Trypanosoma* infection prevalence (Montoya-Porras et al., 2018). In addition, metagenomic studies have shown that in *R. prolixus*, several gut-associated bacteria other than *Rhodococcus* possess the metabolic pathways required for B-vitamin biosynthesis (Tobias et al., 2020). Furthermore, whole-genome sequencing of hosts and their associated symbionts has revealed that *Wolbachia* symbionts are frequently associated with diverse *Rhodnius* species (Filee et al., 2022). Although these *Wolbachia* symbionts display variation in their putative capabilities, this study supports the idea that dual or even multiple nutritional co-symbioses may be common within the Triatominae subfamily. However, the precise relationships and metabolic functions linking the different symbionts and their hosts remain largely unresolved, owing to the limited availability of triatominae (meta)genomics data.

Here, we test the hypothesis that triatominae nutritional symbioses involve a limited set of recurrent bacterial lineages forming stable yet potentially complementary associations within hosts. To address this question, we screened all publicly available Triatominae (meta)genomes (n=18), including 14 generated by our group, to identify bacterial symbionts and detect lateral gene transfers (LGTs). We assembled 15 bacterial symbiont genomes belonging to four genera and investigated their diversity, evolutionary origins, and metabolic potential, revealing patterns of co-infections and metabolic complementarity. We also explore signatures of host–symbiont associations through LGT analyses. Together, our results provide new insights into the complexity and evolution of nutritional symbioses in this major group of insect disease vectors.

## Materials and methods

### Sampling, metagenomes sequencing and assembling

At the time of this study, good quality genome assemblies made from long sequencing reads were available for only four triatominae species, including three from *Triatoma* (*T. dimidiata, T. infestans* and *T. sanguisuga*) and one from a close relative to the *Triatoma* species: *Panstrongylus geniculatus*. We decided to sequence 14 new genomes covering the entire *Rhodnius* phylogeny and add one new *Triatoma* using a combination of short and long sequencing reads (Table 1).

**Table 1:**
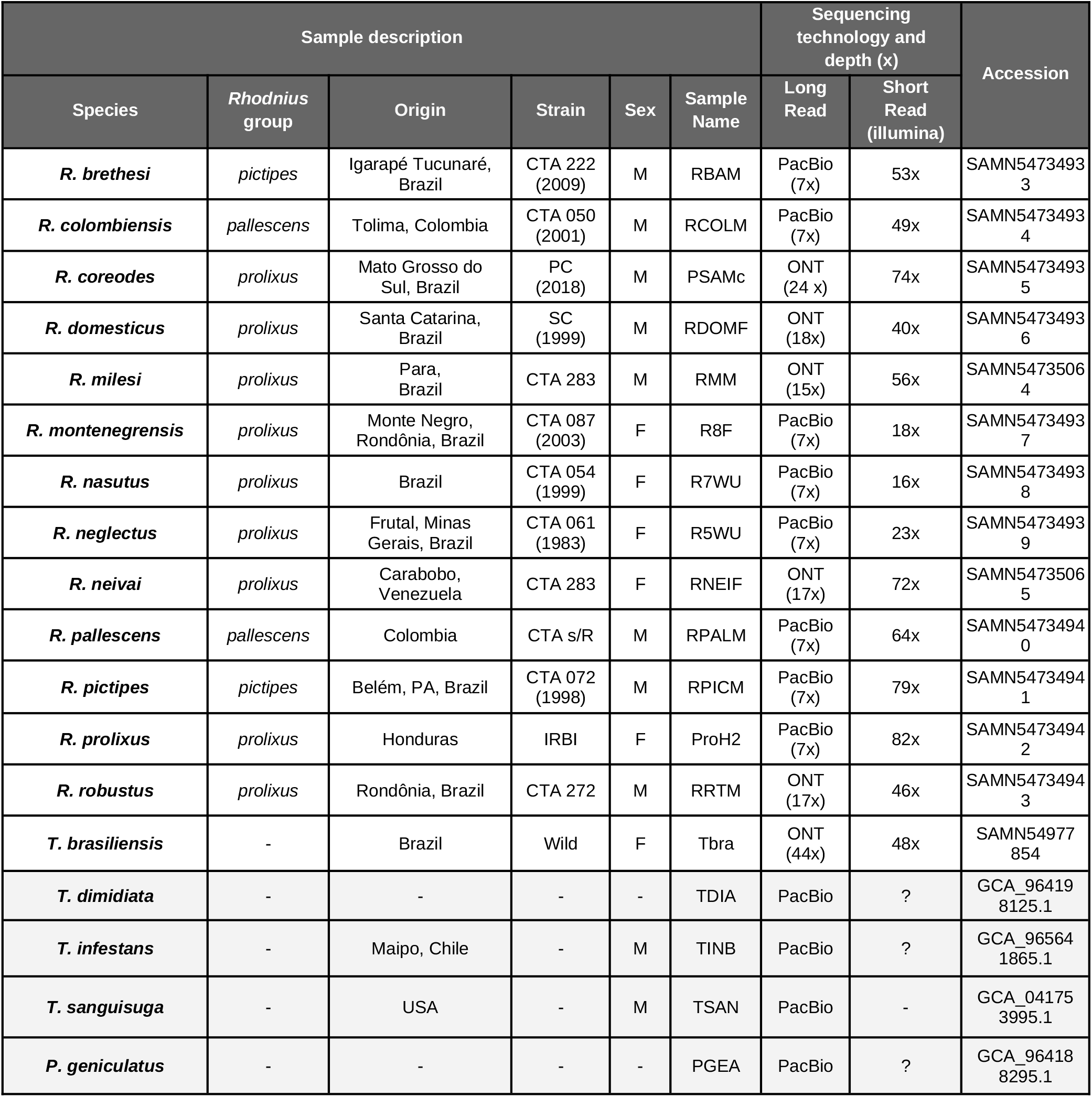
Samples and Sequencing data of studied insects. Genome assemblies already published are indicated with a gray background.

| Sample description |  |  |  |  |  | Sequencing technology and depth (x) |  | Accession |
| --- | --- | --- | --- | --- | --- | --- | --- | --- |
| Species | *Rhodnius* group | Origin | Strain | Sex | Sample Name | Long Read | Short Read (illumina) |  |
| *R. brethesi* | *pictipes* | Igarapé Tucunaré, Brazil | CTA 222 (2009) | M | RBAM | PacBio (7x) | 53x | SAMN54734933 |
| *R. colombiensis* | *pallescent* | Tolima, Colombia | CTA 050 (2001) | M | RCOLM | PacBio (7x) | 49x | SAMN54734934 |
| *R. coreodes* | *prolixus* | Mato Grosso do Sul, Brazil | PC (2018) | M | PSAMc | ONT (24 x) | 74x | SAMN54734935 |
| *R. domesticus* | *prolixus* | Santa Catarina, Brazil | SC (1999) | M | RDOMF | ONT (18x) | 40x | SAMN54734936 |
| *R. milesi* | *prolixus* | Para, Brazil | CTA 283 | M | RMM | ONT (15x) | 56x | SAMN54735064 |
| *R. montenegrensis* | *prolixus* | Monte Negro, Rondônia, Brazil | CTA 087 (2003) | F | R8F | PacBio (7x) | 18x | SAMN54734937 |
| *R. nasutus* | *prolixus* | Brazil | CTA 054 (1999) | F | R7WU | PacBio (7x) | 16x | SAMN54734938 |
| *R. neglectus* | *prolixus* | Frutal, Minas Gerais, Brazil | CTA 061 (1983) | F | R5WU | PacBio (7x) | 23x | SAMN54734939 |
| *R. neivai* | *prolixus* | Carabobo, Venezuela | CTA 283 | F | RNEIF | ONT (17x) | 72x | SAMN54735065 |
| *R. pallescens* | *pallescent* | Colombia | CTA s/R | M | RPALM | PacBio (7x) | 64x | SAMN54734940 |
| *R. pictipes* | *pictipes* | Belém, PA, Brazil | CTA 072 (1998) | M | RPICM | PacBio (7x) | 79x | SAMN54734941 |
| *R. prolixus* | *prolixus* | Honduras | IRBI | F | ProH2 | PacBio (7x) | 82x | SAMN54734942 |
| *R. robustus* | *prolixus* | Rondônia, Brazil | CTA 272 | M | RRTM | ONT (17x) | 46x | SAMN54734943 |
| *T. brasiliensis* | - | Brazil | Wild | F | Tbra | ONT (44x) | 48x | SAMN54977854 |
| *T. dimidiata* | - | - | - | - | TDIA | PacBio | ? | GCA_964198125.1 |
| *T. infestans* | - | Maipo, Chile | - | M | TINB | PacBio | ? | GCA_965641865.1 |
| *T. sanguisuga* | - | USA | - | M | TSAN | PacBio | - | GCA_041753995.1 |
| *P. geniculatus* | - | - | - | - | PGEA | PacBio | ? | GCA_964188295.1 |

We used laboratory CTA strains from the “Insetário de Triatominae da Faculdade de Ciências Farmacêuticas”/Unesp/Araraquara, Brazil (gift from A. Da Rosa & J. Olivieira) and from IRBI lab, France (gift from C. Lazzari) in addition with wild-caughts strains (Table 1). Samples were stored in absolute EtOH except for samples sequenced with the Minion technology stored at -80°C.

DNA was extracted from legs and wing muscles from one individual using the Nucleobond AXG kit (Macherey Nagel). In order to obtain short read sequences (paired-end, 2 x 150 bp), whole-genome shotgun sequencing was performed using Illumina technology, performed with NovaSeq6000-A00318, 150 bp paired-end (Genotoul platform, Toulouse). For three species *R. montenegrensis, R. nasutus* and *R. neglectus* Illumina data were sequenced in a previous study (Filée et al., 2022).

Long read sequences were obtained over a long period of time for which sequencing techniques have improved rapidly. As a consequence, the dataset was obtained with different technologies and coverage. We used Pacbio technology, at the I2BC (Gif-sur-Yvette, France) for 8 species and with MinION technology, performed with GridION GXB02039 at the Genotoul platform (Toulouse, France) for the remaining 6 species.

We performed a *de novo* hybrid assembly for each species using both long reads and shorts reads data using the MaSuRCA v.4 program (Zimin et al., 2013) with the modified version of the CABOG assembler (Miller et al., 2008). To assemble *R. montenegrensis, R. nasutus* and *R. neglectus* genomes, Illumina’s adaptors were removed using Cutadapt v.1.18 (Martin, 2011), and PacBio contigs were auto-corrected using the mode correct of Canu v.1.7 (Koren et al., 2017). To assemble the other species, 150 pb Illumina reads and GridIon reads were used without any upstream processing.

### Identification of Symbiont Genomes

Bacterial contigs were identified in host genome assemblies based on a combination of sequence length, GC content, read coverage, and taxonomic assignment using BlobTools (Laetsch and Blaxter, 2017). Read coverage was estimated by mapping raw reads to each contig using minimap2 (Li, 2018). BlobTools also provided an initial bacterial taxonomic assignment by annotating contigs through DIAMOND searches (Buchfink et al., 2015) against the UniProt UniRef90 database. Contigs assigned to prokaryotes were subsequently annotated with Prokka (Seemann, 2014). Genome completeness and contamination were estimated using CheckM (Parks et al, 2015). The GMX-tools package was used to compute assembly statistics such as N50 and N90. Genome circularity of each bacterial assembly was assessed by mapping the Oxford Nanopore (MinION) reads to the end of contigs using minimap2 (Li, 2018). Alignments were visualised in IGV (Robinson et al., 2011) to determine whether reads spanned both contig ends. For each symbiont genus, gene presence-absence patterns were determined using BLASTP (e-value< 1e-5) allowing the identification of conserved and missing genomic regions across symbiont genomes.

### Lateral Gene Transfer (LGT) analysis

Contigs displaying read coverage and GC content similar to the host genome but showing bacterial sequence similarities were extracted based on BlobTools results. To exclude residual bacterial contigs a conservative threshold was applied, *i*.*e*., contigs for which more than 65% of their total length matched bacterial genes were considered likely bacterial in origin and removed as false positives. Another taxonomic assignment was performed using DIAMOND searches against the UniProt database, to filter contigs that did not harbour at least one eukaryotic gene. After this step, all the remaining contigs are thus composed of a mix of eukaryotic and bacterial-like genes. Genes from Triatominae-associated symbionts identified in this study were then queried against all the contigs that have successfully passed the previous filter. Gene hits were retained if they showed an e-value < 1e-5 and > 99% sequence identity with a symbiont gene sequence. Loci matching multiple bacterial genera were classified as “unassigned”. These stringent criteria were used to have reliable and unambiguous taxonomic assignments.

### Phylogeny of Symbiont Genomes

A database of bacterial genomes was compiled from NCBI, including reference genomes for each of the four genera identified in this study (Supplementary Table 1). Genomes with uncertain origins or host associations were excluded. Orthologous genes were identified using OrthoFinder. Orthologous sequences were then aligned with MAFFT (Katoh et al., 2002) using the L-INS-i algorithm and concatenated into a supermatrix. Maximum-likelihood phylogenies were inferred with IQ-TREE 2.2.2.6 (Minh et al., 2020), with 1,000 ultrafast bootstrap replicates. The best-fitting substitution model was selected using ModelFinder (Kalyaanamoorthy et al., 2017).

### Metabolic annotation

Key metabolic pathways were initially identified by querying the annotated amino acid sequences against BlastKOALA (Kanehisa et al., 2016). B-vitamin biosynthesis and amino acid production pathways were then examined in detail by building a custom database of (Grant et al., 2023) homologous genes from diverse bacterial insect symbionts (Supplementary Table 2). These reference proteins were used as queries in BLASTP searches (Altschul et al., 1990) against the symbiont genome assemblies, retaining matches with an e-value below 1e-5.

### Gene conservation analysis

To detect the pattern of gene presence/absence among symbiont genomes we used BLASTP (Altschul et al., 1990): each ORFs from a given genome was blasted against the ORFs of the other genomes retaining hits with an e-value cutoff of 1e-5. The resulting tabular output files were then analyzed by the Proksee software (Grant et al., 2023) to generate the graphical maps.

### Genome alignment

Homologous genes between the two *Arsenophonus* symbiont genomes identified in *R. domesticus* were compared using BLASTN (Altschul et al., 1990) to assess gene presence-absence patterns and detect putative deletions. To localize deletions events, genes from the smaller genome were aligned against the homologous genes present in the largest *Arsenophonus* assembly. Genes displaying deletions were subsequently analyzed with BlastKOALA (Kanehisa et al., 2016) to infer their putative functional roles.

## Results

### Symbiont genomes assemblies and annotation

We used BlobTools graphs to discriminate host and symbiont contigs based on GC content, read coverage and sequence similarity (Figure 1). In nine genome assemblies distinct contigs deviated from the insect contig distribution, indicating the presence of symbiont sequences. For example in the *R. brethesi* and *R. robustus* genome assemblies, there are two different symbionts: in *R. brethesi* a *Wolbachia* (orange spots) and a *Rhodococcus* (purple spots), in *R. robustus*, a *Symbiopectobacterium* (orange spots) and a *Rhodococcus* (purple spots) (Figure 1). In total, 15 different potential symbiont genomes were identified (Table 2 and Supplementary Figure 1), and co-infection by at least two symbionts seems frequent (5 cases). The annotation of these contigs indicated that these symbionts belong to four genera: *Arsenophonus, Rhodococcus, Symbiopectobacterium* and *Wolbachia*.

**Figure 1:**
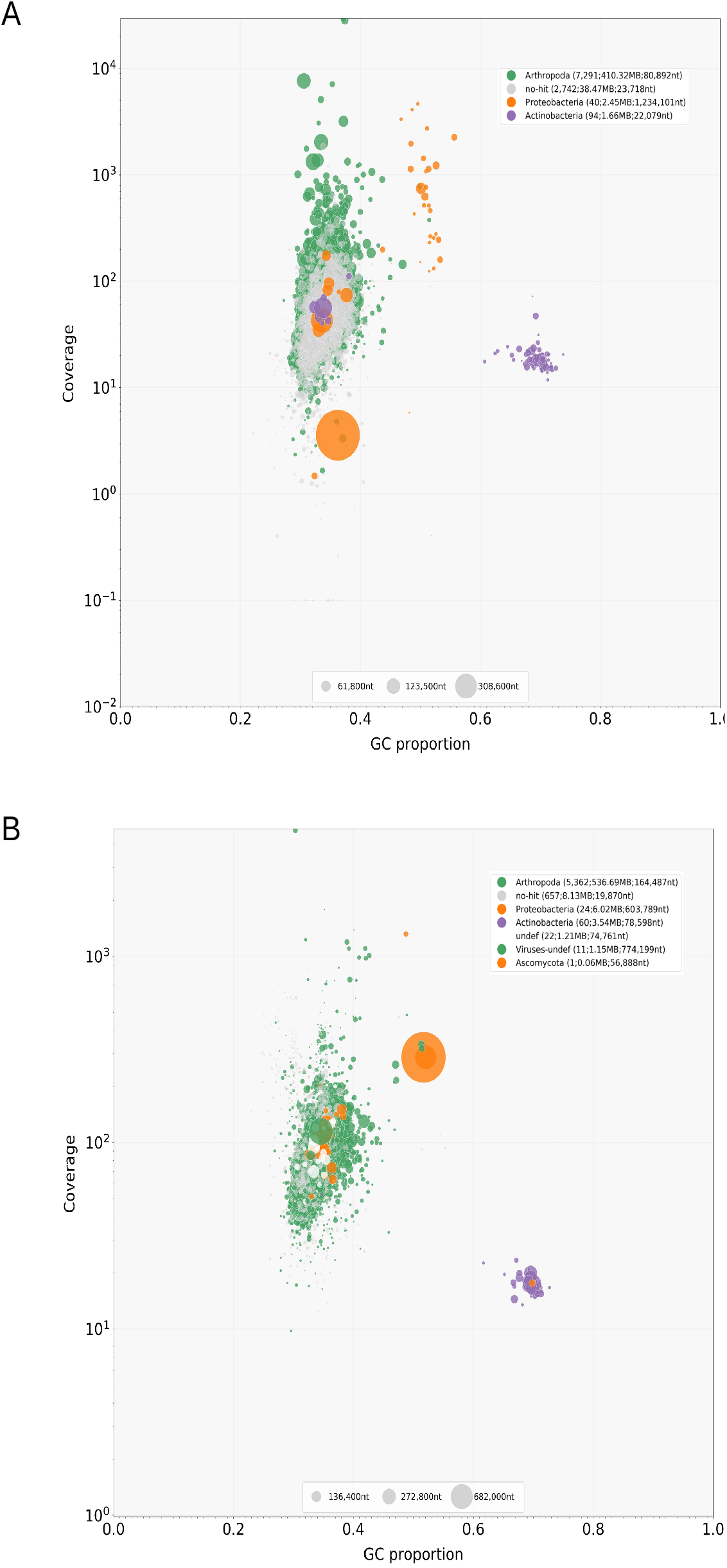
Detection of symbionts within the metagenomes of *R. brethesi* (A) and *R. robustus* (B). Contigs represented as circles were binned according to their GC%, read coverage, and taxonomic assignment, with circle size corresponding to contig length.

**Table 2:** Bacterial genomes characteristics. Rows are colored based on the taxonomic assignment. For genomes assembled as a single contig, circularity was evaluated (yes: confirmed)

| Host | Accession | Contigs (#) | Genome size (Mb) | GC content (%) | N50 (Kb) | Completeness (%) | Circularity |
| --- | --- | --- | --- | --- | --- | --- | --- |
| *T. brasiliensis* (Tbra) | *Arsenophonus* | 20 | 3.78 | 38.5 | 404.26 | 80.54 | - |
| *R. brethesi* (RBAM) | *Rhodococcus* | 89 | 1.36 | 65.2 | 19.62 | 21.54 | - |
|  | *Wolbachia* | 1 | 1.23 | 36.3 | 1 234.10 | 99.15 | yes |
| *R. pictipes* (RPICM) | *Rhodococcus* | 102 | 4.18 | 69.7 | 63.25 | 90.6 | - |
|  | *Wolbachia* | 1 | 1.21 | 36.2 | 1 214.27 | 99.57 | yes |
| *R. coreodes* (PSAMc) | *Symbiopectobacterium* | 1 | 3.65 | 49.8 | 3 647.24 | 84.78 | yes |
| *R. domesticus* (RDOMFM) | *Arsenophonus* | 1 | 1.15 | 38.2 | 1 151.04 | 29.73 | yes |
|  | *Arsenophonus* | 5 | 3.47 | 37.8 | 744.93 | 76.79 | - |
|  | *Symbiopectobacterium* | 5 | 3.71 | 50.1 | 2 735.30 | 78.11 | - |
| *R. milesi* (RMM) | *Rhodococcus* | 4 | 4.68 | 69.4 | 4 541.84 | 97.12 | - |
|  | *Symbiopectobacterium* | 11 | 3.75 | 51.7 | 1 214.85 | 72.12 | - |
| *R. neglectus* (R5WU) | *Wolbachia* | 8 | 0.98 | 36.3 | 978.41 | 53.85 | - |
| *R. neivai* (RNEIF) | *Symbiopectobacterium* | 10 | 3.32 | 51.3 | 1 130.79 | 51.72 | - |
| *R. robustus* (RRTM) | *Rhodococcus* | 59 | 3.53 | 69.5 | 78.60 | 72.08 | - |
|  | *Symbiopectobacterium* | 6 | 5.13 | 51.6 | 2 727.91 | 92.56 | - |

The presence of a *Rhodococcus* symbiont was systematically associated with an additional symbiont either a *Symbiopectobacterium* or a *Wolbachia* suggesting a recurrent pattern of co-occurrence. In *R. domesticus*, two different *Arsenophonus* genome assemblies have been evidenced: a small assembly (single contig, ∼ 1 Mb), and a large one (five contigs, ∼ 3.5 Mb). Despite its small size, the smallest *Arsenophonus* genome assembly was circularized, as several long reads map to both the 5’ and 3’ ends of the contig. Genome size and completeness variations were also noticed for the other symbiont genus (Table 2). For instance, with 5.13 Mb, the *Symbiopectobacterium* genome found in *R. robustus* was about 1.54 to 1.36 larger than those associated with the other hosts. Even circularized, single-contig assemblies display lower completeness such as the *Symbiopectobacterium* associated with *R. coreodes* (85% completeness). These patterns are consistent with varying degrees of genome reduction among symbionts, although differences in sequencing and assembly approaches may also contribute to the observed variation. All single-contig genomes were confirmed to be circular. Differences in sequencing technologies and genome assembly pipelines may influence the detection of symbiont-derived sequences, particularly in publicly available genomes. In some cases, bacterial contigs may have been removed during host genome curation steps. Therefore, the absence of detectable symbiont sequences in some assemblies should be interpreted cautiously.

### Lateral Gene Transfer Analysis

BlobTools analyses revealed typical host contigs with bacterial taxonomic similarity despite host-like coverage and GC content (Figure 1). To validate this observation, we mapped symbiont sequences against the host genomes. We applied stringent criteria, retaining only contigs that display a combination of eukaryotic-like and bacterial-like gene signature to avoid misclassification of free bacterial symbionts as LGTs. This analysis revealed putative lateral gene transfers (LGTs) between symbionts and hosts (Figure 2). Twelve out of 13 *Rhodnius* genomes harbour *Wolbachia* genes, even when no *Wolbachia* genome was retrieved from the assembly. *Wolbachia* has not been detected in *Triatoma/Panstrongylus* species. LGT with *Arsenophonus* genes were only detected in the three host genomes for which *Arsenophonus* symbiont genomes were retrieved from the assembly (*R. domesticus, T. brasiliensis* and *P. geniculatus*). In contrast *Symbiopectobacterium* genes were found in all host genomes for which a *Symbiopectobacterium* symbiont genome was retrieved from the assembly, as well as in two additional host genomes. In *R. milesi*, these genes were found on the same contig ( ∼ 70kb) covering fourth of the total contig length. Finally, *Rhodococcus* sequences were present in all *Triatoma* and *Panstrongylus* species and displayed a more scattered distribution in *Rhodnius* genomes, particularly when a *Rhodococcus* symbiont was detected in the assembly. Putative functions of the laterally transferred genes are very diverse with no clear bias towards specific pathways (Supplementary Figure 2)

On average, each triatominae genome harbored genetic traces of at least two symbiont genera and six host genomes from three or more. *R. domesticus* displayed symbiont genes from the four symbiont genera. The number of laterally acquired highly varied across hosts with some host genomes containing more than 100 genes highly similar to symbiont genes (Figure 2). Notably, the presence of laterally transferred genes was strongly associated with the detection of the corresponding symbiont genome. Conversely, all triatominae species with no detectable free-living symbiont in assemblies also harbored signatures of LGTs from various symbiont origins. This is especially true for *Wolbachia* in *Rhodnius species* and *Rhodococcus* in *Triatoma/Panstrongylus* species for which LGTs were widespread. Lateral gene transfers from the symbionts can thus be used as “footprint” of present or past association, even for insect hosts for which genome assemblies do not reveal any symbionts. In this way, our data reveals that most of our triatominae species we have analyzed are (or have been) associated by the 4 different symbionts.

**Figure 2:**
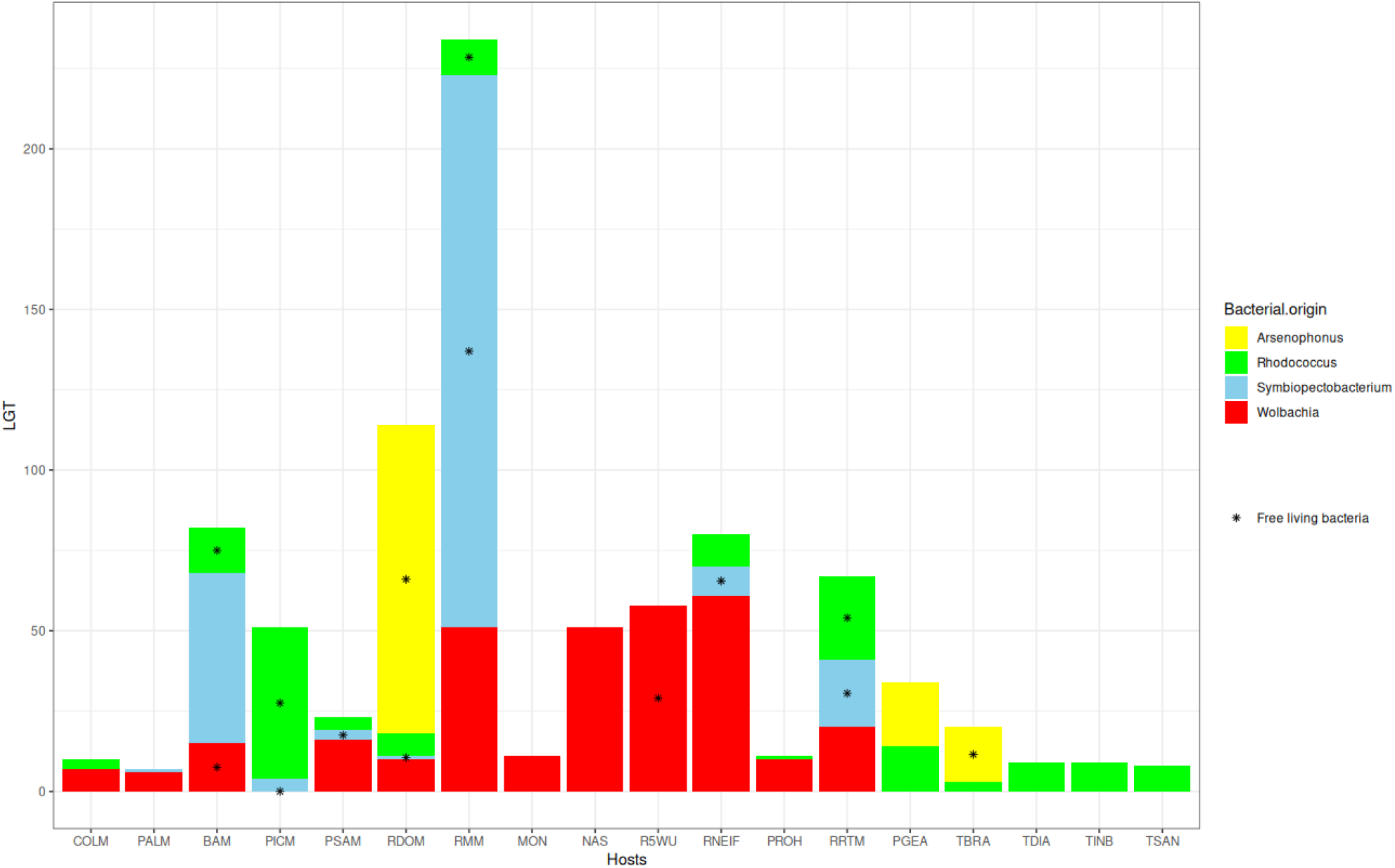
Amount of symbiont genes in the triatominae host genomes. Each bar represents the number of laterally transferred genes (LGT) per host genomes. Bars are colored according to the symbiont genus. A black dot indicates that a corresponding symbiont genome was identified and assembled for this insect.

### Symbiont Genome Phylogenies

To better understand the evolutionary origins of the symbionts, we analysed the phylogenetic relationships of the four symbiont genera present in Triatominae based on a set of single-copy orthologous genes (Figure 3). For each genus, Triatominae-associated symbionts formed a monophyletic group with branches generally short and displaying low internal resolution. These results support a single origin for each symbiont present in Triatominae, even for symbionts infecting distantly related Triatominae hosts, such as *Arsenophonus* associated with *T. brasiliensis* and several *Rhodnius* species (host divergence time >20 My ; Hernández et al., 2025. In this phylogeny (Figure 3a), the largest *Arsenophonus* genome assembly is closer to the other triatominae symbionts than the smaller one, confirming that two different *Arsenophonus* symbionts with a divergent evolutionary history co-exist in the same host. Except for *Rhodococcus*, the three other triatominae-associated symbionts clustered with other insect symbionts, including hematophagous hosts. For instance, triatominae-associated *Arsenophonus* form a monophyletic group with Hippoboscidae louse fly symbionts, and triatominae-associated *Wolbachia* are placed within a group composed of either symbionts of other hematophagous insects (the bedbug *Cimex lectularius*) or symbionts of organisms with an hemoparasitic lifecycle (mainly nematodes). *Symbiopectobacterium* symbionts appear to be related to the symbiont of another Heteroptera: the granivorous bulrush bug *Chilacis typhae*. All of these related symbionts are known (or are supposed) to be nutritional symbionts of their hosts, suggesting an ancestral adaptation of these groups to nutritional symbioses associated with nutrient-poor feeding (blood or plant-based). Finally, *Rhodococcus* associated with triatominae displayed a different pattern, the closest matches correspond to telluric, free-living, bacteria from soils, highlighting the fact that, unlike *Arsenophonus, Symbiopectobacterium, and Wolbachia*, that are primarily endosymbiotic, *Rhodococcus* is mostly free-living saprotrophic bacteria.

**Figure 3:**
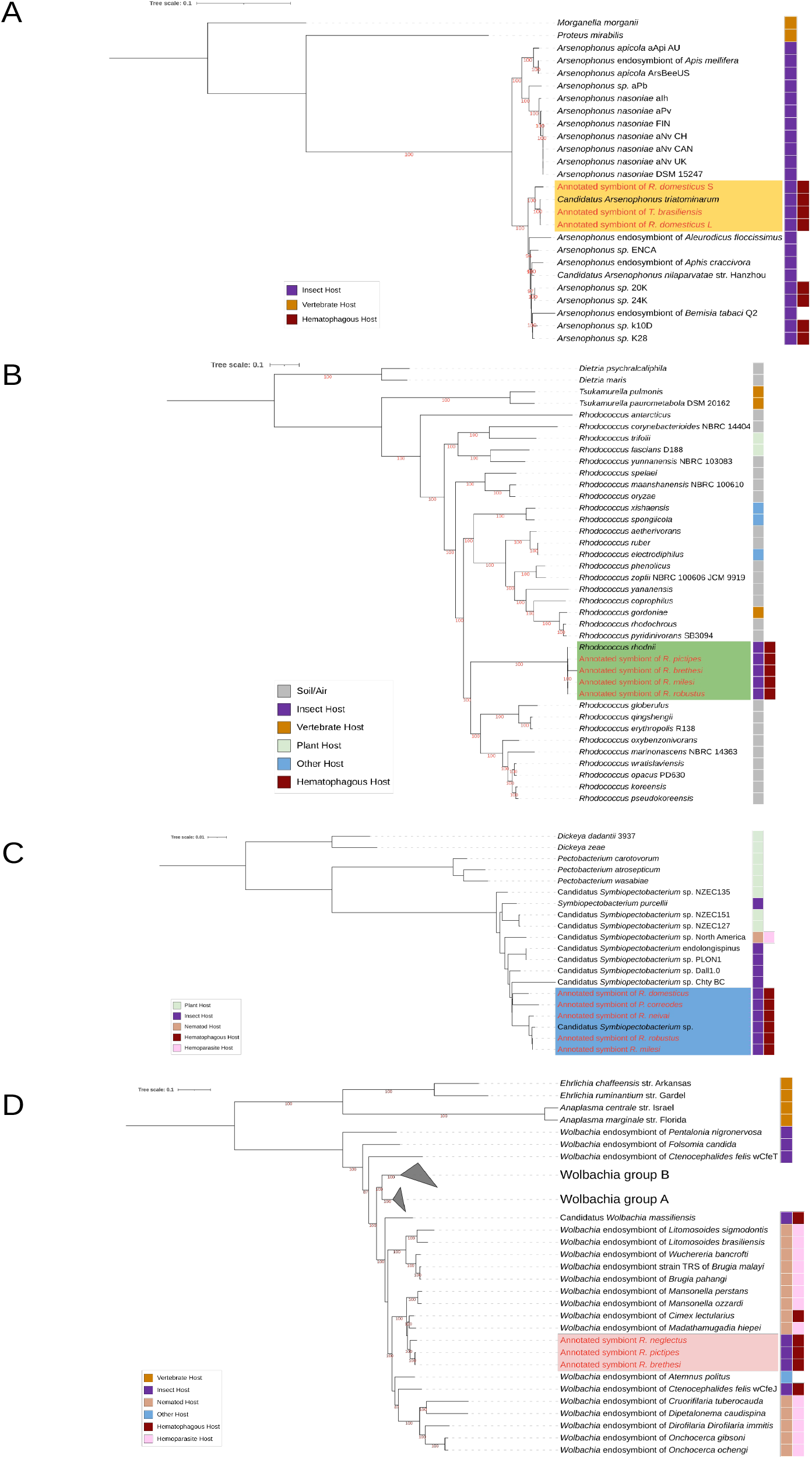
Phylogenies of triatominae symbionts. Each tree represents the best maximum-likelihood tree inferred from conserved single-copy orthologs. Ultrafast bootstrap values (1,000 replicates) above 80 are indicated in red below the corresponding node. Colored ranges indicate the clades formed by Triatominae-associated symbionts. **A:** *Arsenophonus* tree based on 33 orthologs (9,047 aa). **B:** *Rhodococcus* tree based on 117 orthologs (38,473 aa). **C:** *Symbiopectobacterium* tree based on 178 orthologs (58,772 aa). **D:** *Wolbachia* tree based on 44 orthologs (13,124 aa). The fully expanded *Wolbachia* phylogeny is provided in Supplementary Figure 3.

### Gene conservation analysis

Patterns of gene presence/absence across all four genera revealed that most genes are highly conserved among the different symbionts, confirming the close evolutionary relationship of Triatominae-associated symbionts (Figure 4). Discrepancies between symbiont genomes were not localized to specific regions but were distributed throughout the genome. This pattern supports the notion that genome erosion occurred largely at random, although technical limitations in sequencing and assembly may also contribute to the observed variation. Regarding the two *Arsenophonus* genomes associated with *R. domesticus*, 337 shared genes exhibited alignments that were more than 30% shorter in the smaller genome (Figure 5). These deletions appeared to be randomly distributed along the alignments, with no evidence for specific genomic regions being preferentially targeted by sequence reduction. A comparative analysis of metabolic capabilities between the two *Arsenophonus* symbionts revealed that genes present in the larger genome but absent from the smaller one predominantly belonged to three functional categories: “Unclassified genetic information processing”, “Energy metabolism” and “carbohydrate metabolism” (Supplementary Figure 4). This analysis supports the hypothesis that the smaller *Arsenophonus* genome results from an ongoing process of genome erosion affecting most regions of the genome, with a more pronounced impact on several metabolic pathways.

**Figure 4:**
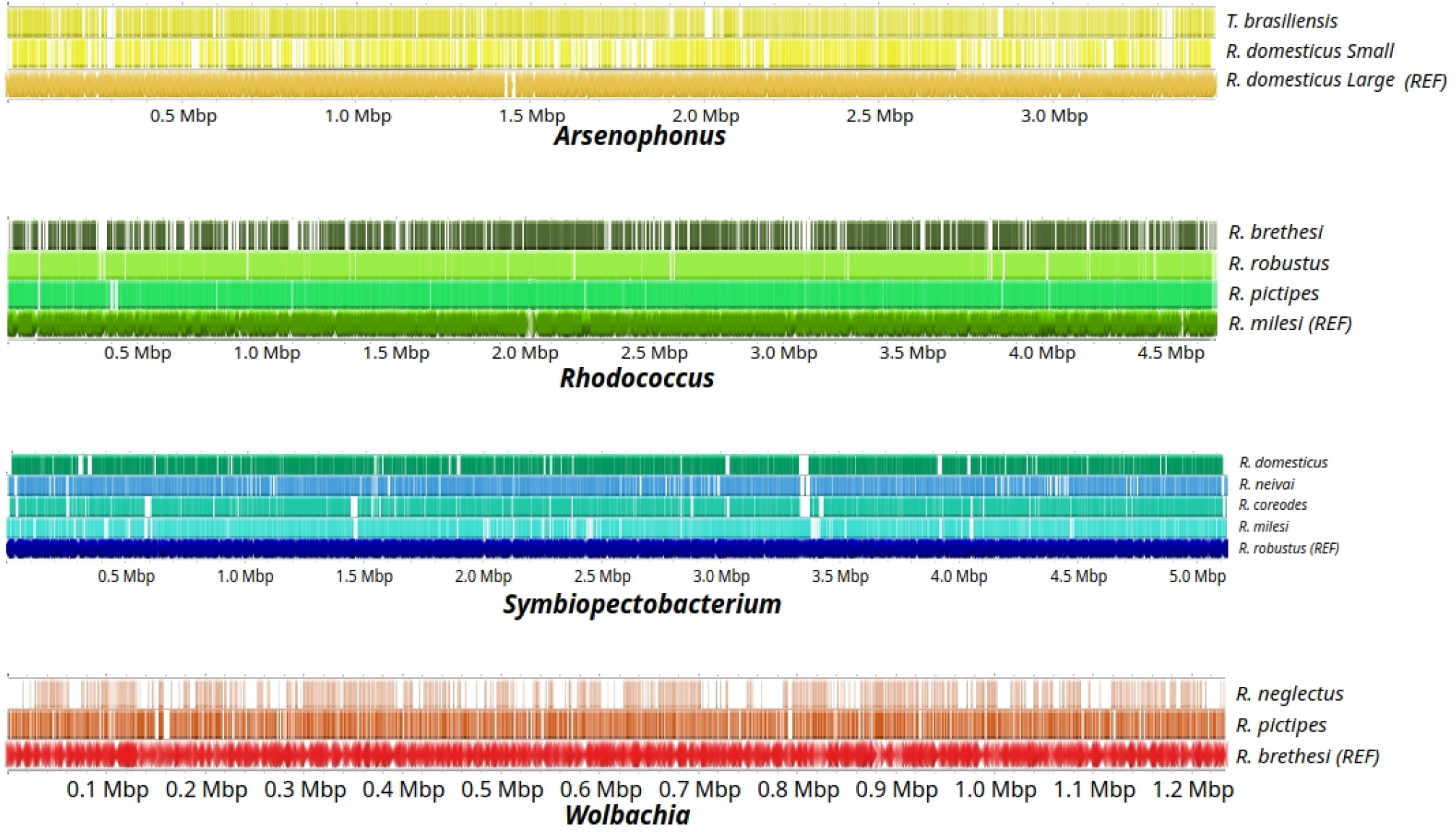
Gene conservation for each symbiont genus. The arrows in the lower lanes of each symbiont panel correspond to the position of the different ORFs present in the reference genomes annotated as “REF”. Traits represent the presence of the ORFs in the different corresponding genomes.

**Figure 5:**
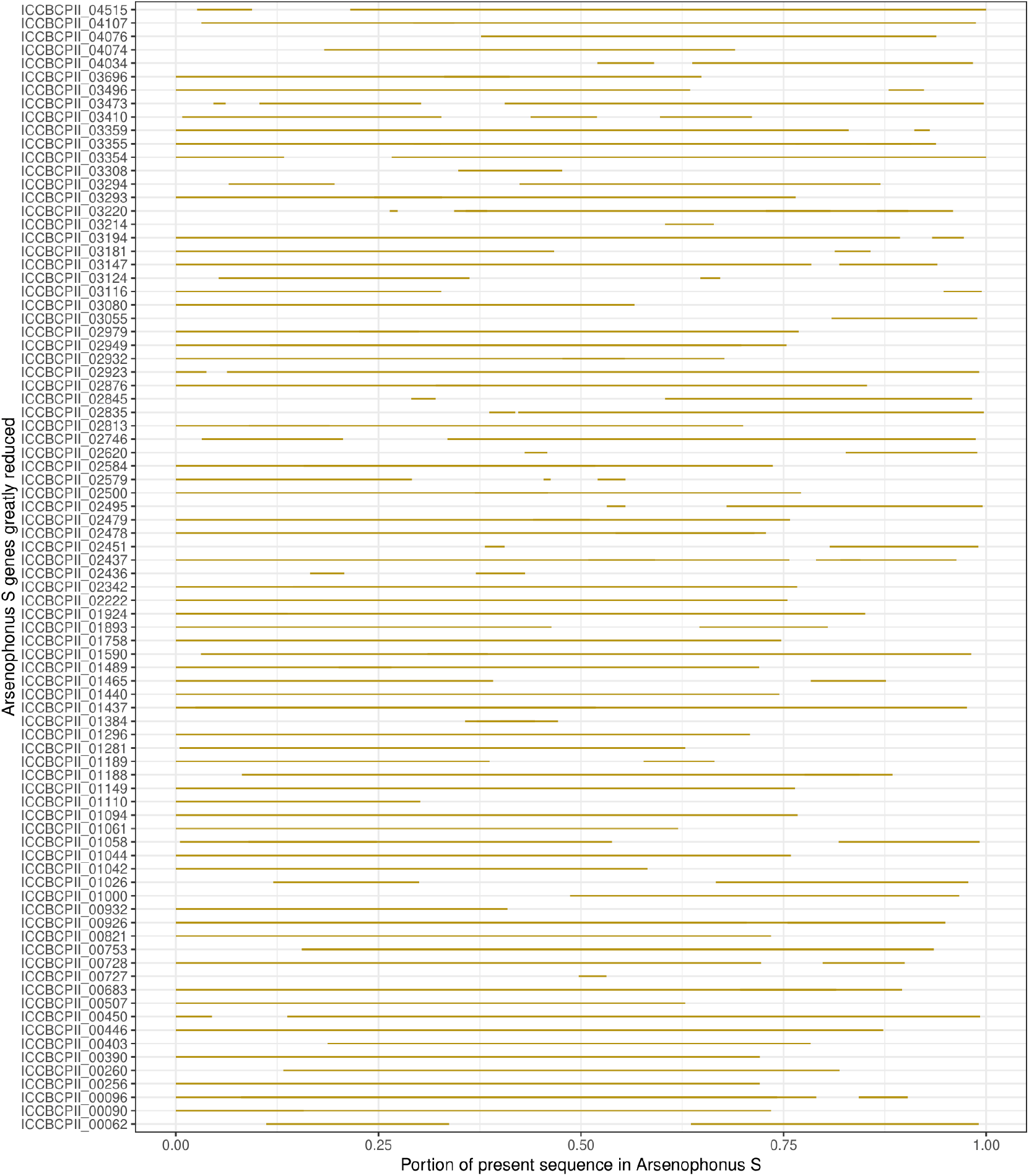
Alignments of 100 homologous ORFs shared by the two *Arsenophonus* symbiont genomes associated with *R. domesticus*. Genes from the smaller assembly were aligned against their homologs in the larger assembly. Each yellow segment indicates the genomic portion of each homologous gene sequence present in both genomes.

### Metabolic gene annotation and comparison

To better characterize the metabolic potential of each symbiont, we assessed the completeness of key metabolic pathways (Figure 6). With few exceptions, most symbionts exhibited complete metabolic pathways for amino acid biosynthesis, supporting the notion that they do not rely entirely on the host for amino acid provisioning. Notable exceptions included the lack of several folate-related genes in all *Rhodococcus* genomes and incomplete cobalamin (vitamin B12) pathways in both *Wolbachia* and *Rhodococcus* genomes.

**Figure 6:**
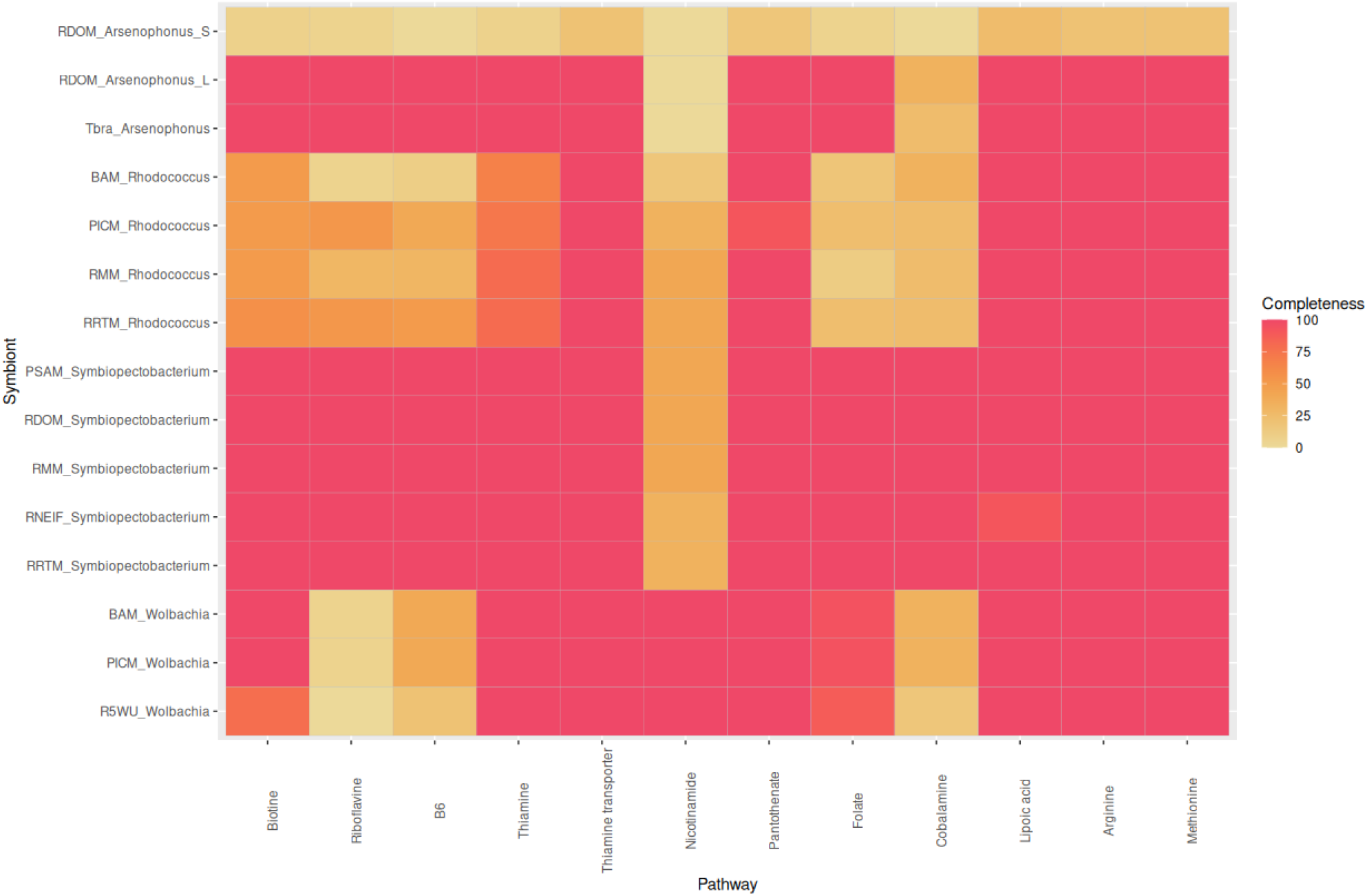
Metabolic pathways in Triatominae symbionts. Symbionts are ordered by bacterial genus. Pathway completeness corresponds to the percentage of genes involved in each metabolic pathway that were annotated in the symbiont genome assembly.

None of the symbionts harbored fully complete B-vitamin biosynthesis pathways, suggesting varying degrees of metabolic specialization. *Symbiopectobacterium* and the two largest *Arsenophonus* genomes displayed the most complete sets of B-vitamin pathways, lacking only the nicotinamide (vitamin B3) pathway (Figure 6). In general, symbionts belonging to the same bacterial genus showed similar patterns of gene presence/absence. *Wolbachia* genomes were characterized by near-complete loss of riboflavin (vitamin B2) and vitamin B6 pathways. Likewise, all *Rhodococcus* symbionts exhibited multiple incomplete B-vitamin pathways, including those involved in biotin (vitamin B7), riboflavin (vitamin B2) and vitamin B6 biosynthesis. However, *Rhodococcus* symbionts were consistently found in association with either *Symbiopectobacterium* or *Wolbachia*, which retain the capacity to complement some or all of these pathways. For instance, the *Rhodococcus* genomes associated with *R. robustus* and *R. milesi* lacked most B-vitamin biosynthesis genes, whereas their associated *Symbiopectobacterium* symbionts possessed the metabolic potential to compensate for these deficiencies. Similar patterns of metabolic complementarity were observed between *Rhodococcus* and *Wolbachia* co-occurring in *R. brethesi* and *R. pictipes*. Together, these results support the existence of possible B-vitamin metabolic supplementation by the symbionts but also some degree of complementation among co-infecting symbionts within the same host.

In contrast, the smaller *Arsenophonus* genome identified in *R. domesticus* displayed a markedly different profile: all examined metabolic pathways were either incomplete or entirely absent, including amino acid biosynthesis pathways that were otherwise well conserved in other symbionts. This finding supports our previous analyses indicating extensive genome erosion in this symbiont and suggests that it is fully dependent on metabolic complementation from other conspecific symbionts and/or the host for its development.

## Discussion

More than 90 years ago, Wigglesworth demonstrated that *R. prolixus* critically depends on a gut bacterial symbiont, historically assigned to the genus *Rhodococcus* (Wigglesworth, 1930). This observation led to the hypothesis of a nutritional symbiosis supplementing the nutrient-poor blood diet of triatominae. However, the identity of the symbionts involved and the specific metabolites supporting the symbiosis have remained unclear. Our metagenomic analysis of 18 triatominae genomes, including 13 *Rhodnius* species and 5 *Triatoma/Panstrongylus* species, aimed to identify the symbionts, characterize their metabolic potential, and explore the evolutionary dynamics of these associations.

### Multiple symbionts mediate nutritional symbiosis

In eight *Rhodnius* species our data suggest that nutritional symbiosis involves four bacterial lineages: *Rhodococcus, Arsenophonus, Symbiopectobacterium*, and *Wolbachia*. We retrieved four *Rhodococcus* genomes across four *Rhodnius* species, supporting a central role for this taxon in the symbiotic system of Triatominae. Notably, the presence of a *Rhodococcus* genome was consistently associated with the co-occurence of at least one additional genome, either *Wolbachia* or *Symbiopectobacterium*. In addition, we recovered three *Arsenophonus* genomes, detected either as single symbionts or in association with *Symbiopectobacterium*.

Taken together, these results support the hypothesis that nutritional symbiosis in Triatominae is not mediated by a single obligate symbiont, but rather by a multipartite symbiotic system. This organization may promote metabolic complementarity, functional redundancy, or increased resilience to symbiont loss. Consistent with this hypothesis, all three additional symbiont lineages identified in this study are phylogenetically closely related to well-characterized obligate and/or nutritionally important insect symbionts. *Wolbachia* strains recovered in this study cluster with the nutritional *Wolbachia* symbiont of bedbugs, which has been shown to provide essential B vitamins to its hosts (Nikoh et al., 2014). Similarly, *Symbiopectobacterium* symbionts are phylogenetically closely related to obligate nutritional symbionts described in a wide range of insects and nematodes where they contribute to host metabolism and development (Martinson et al., 2020).

Finally, *Arsenophonus* symbiont are related to lineages commonly found in plant-based and blood-feeding insect, many of which are strongly suspected or experimentally demonstrated, to function as nutritional providers (Martin Říhová et al., 2023). More broadly, insects belonging to the order Hemiptera appear particularly prone to hosting multiple nutritional symbionts. This pattern has been extensively documented in sap-feeding hemipterans, where complex symbiotic associations are the rule rather than the exception (Bennett and Moran, 2013; Kobiałka et al., 2018; Manzano-Marín et al., 2023; Szabó et al., 2022), and has also been reported in blood-feeding hemipterans such bedbugs (Hypša et al., 2025). Despite accumulating genomic evidence, the functional contributions and reciprocal benefits of each symbiont within these symbiont associations remain poorly understood. Genomic predictions alone are insufficient and must be complemented by experimental approaches, such as the rearing of axenic insects followed by controlled reinfection with single or multiple symbionts, as well as targeted metabolic assays, to disentangle metabolic dependencies and symbiont interactions.

### Metabolic Complementation and/or tissue specialization

Comparative analysis of the metabolic repertoires of the symbionts examined in this study reveals clear patterns of metabolic complementarity among co-occurring symbionts. Strikingly, all four *Rhodococcus* symbiont genomes encode for incomplete B-vitamin biosynthesis pathways, despite the central role of B vitamins in insect nutritional symbioses. In particular, the *Rhodococcus* genomes recovered from triatominae lack the upstream genes required for the synthesis of the pimeloyl-CoA intermediate (notably f*abB, fabF, fabZ, fabI* and *bioH* genes), which is essential for biotin (vitamin B7) production. In contrast, the downstream biotin pathway genes (*bioA, bioB, bioC, bioD, bioF*) are conserved. This suggests that *Rhodococcus* symbionts may not be able to synthesize biotin de novo and could rely on external sources of precursors, either from the host or from co-infecting symbionts. A similar pattern exists for the riboflavin (B2 vitamin) pathway, which is highly reduced in these *Rhodococcus* genomes. Interestingly, *Rhodococcus* symbionts are consistently associated with either *Symbiopectobacterium* or *Wolbachia*, both of which encode proteins that can fully or partially complement the B-vitamin biosynthesis deficiencies. These observations suggest B-vitamin provisioning by symbionts and highlight potential metabolic complementarity within co-infecting communities. In addition, differences in tissue tropism may contribute to symbiont coexistence. While *Rhodococcus* is known to primarily inhabit the digestive tract, *Arsenophonus, Symbiopectobacterium* and *Wolbachia* are typically endosymbionts capable of infecting multiple tissues. This raises the possibility that co-existing symbionts display tissue-specific distribution, as observed in planthoppers (Michalik et al., 2023) and aphids (Renoz et al., 2022), or cell-specific localization as observed in aphids (Yorimoto et al., 2022). Such spatial partitioning may reflect underlying metabolic interactions, whereby symbionts with complementary pathways co-localize, while those with redundant functions are segregated that minimize competition. This pattern may explain the frequent co-occurrence of multiple symbionts observed across the triatominae species examined here. Finally the exact nature of the metabolic interactions remains to be fully resolved. Although B-vitamins have long been considered central to the *Rhodococcus*-*R. prolixus* symbiosis, recent experiments show that dietary supplementation of axenic *R. prolixus* with B-vitamins fails to restore standard host fitness (Gilliland et al., 2023). This suggests that additional metabolites are likely involved, and future experimental work will be required to identify the metabolites provided by each symbiont and validate their functional roles.

### Symbiont genome evolution in Triatominae

The comparison of the two *Arsenophonus* genomes co-occurring in *R. domesticus* provides insight into the evolutionary dynamics of symbiosis. One genome exhibits typical features of genome degradation : a substantial reduction in genome size, an abundance of truncated genes, and complete loss of most metabolic pathways associated with nutritional symbiosis. Such genome reduction is consistent with patterns observed in other *Arsenophonus* genomes including highly streamlined genomes of less than 1 Mb (Siozios et al., 2024), as well as cases *Arsenophonus* co-occurred with another symbiont (Yorimoto et al., 2022). This suggests that symbiont genome erosion may ultimately reduce symbiont fitness to such an extent that symbiont replacement becomes necessary. The co-infection of two *Arsenophonus* strains in *R. domesticus*, with evidence of genome streamlining for one of them, may represent the early stage of such a replacement process. However, symbiont turnover in Triatominae appears constrained, as all identified symbionts belong to monophyletic clades, suggesting a single association event for each symbiont genus. These constraints may reflect strong host genetic prerequisites, implying that only a specific bacteria can establish stable symbiotic relationships. Such constraints may also slow genome streamlining, as most *Arsenophonus, Rhodococcus* and *Symbiopectobacterium* genomes display large genome size with little evidence of degradation.

### Limited symbiont diversity and constrained dynamics

Despite the high diversity of insect symbiotes reported in the literature, Triatominae appear to harbour a restricted set of 4 symbiont lineages that form monophyletic groups in their respective phylogenies, suggesting a single founder effect at the origin of both lineages. In cimicid species (bedbugs), four different symbiont genera have also been identified, corresponding to 23 symbiont genome assemblies. However, phylogenies indicate at least 14 independent symbiont acquisition events and a high rate of symbiont turnover (Hypša et al., 2025). Similarly, the association between louse flies and *Arsenophonus* arose independently at least four times, supporting multiple replacement events (Martin Říhová et al., 2023). Notably, even when considering the traces of past interactions revealed by lateral gene transfers (LGTs), only a limited number of additional past or cryptic symbiotic associations can be inferred. Most of the Triatominae species analyzed here have been associated with at least two, and on average three, of the four bacterial genera identified so far. This includes specimens for which no symbiont assembly was recovered, or whose symbionts differ from the assembled genera. In contrast, the genome of the bedbug *Cimex lectularius* encode numerous symbiont LGTs originating from a broader diversity of symbiont lineages than Triatominae, including *Wolbachia, Arsenophonus, Sodalis, Symbiopectobacterium*, and *Hamiltonella* (Benoit et al., 2016). These data suggest that nutritional symbiosis in Triatominae involves a restricted network of four major symbiont lineages, smaller than that observed in other blood-feeding insects. Reduced symbiont diversity and limited competition may in turn constrain symbiont turnover in triatominae. Accordingly, our results support the view that symbiont dynamics in Chagas’disease vectors are constrained, involving a limited number of symbiont acquisition events and relatively stable associations over evolutionary time.

## Conclusion

Although Triatominae are an ancient and diverse group, our understanding of their nutritional symbiosis remains largely based on studies of *R. prolixus* and both experimental and genomic data are still limited. By analyzing metagenomic data from a broad sampling Triatominae species, we revealed that nutritional symbiosis involves a limited set of four bacterial lineages, each likely originating from a single event of symbiont acquisition followed by co-evolution. Symbiont genomes exhibit limited degradation, although B-vitamin biosynthesis pathways are frequently incomplete. These deficiencies may be compensated by co-infecting symbionts encoding genes with complementary metabolic functions. Our results support a model in which nutritional symbiosis in Triatominae relies on a small but relatively stable network of symbionts, with limited evidence for competition and symbiont replacement. We propose that the gut *Rhodococcus* symbiont, which appears widespread among Triatominae, may be complemented by diverse endosymbionts infecting different tissues or cells, thereby minimizing direct inter-specific competition. These genomic predictions call for experimental validations,focusing not only on the *R. prolixus* species host but also on other Triatominae hosts. Finally, these findings may have applications for vector control strategies. Targeting key symbionts or disrupting essential metabolic pathways could potentially reduce host fitness, thereby providing a basis for the development of symbiont-based control strategies.

## Acknowledgments

We would like to thank all our collaborators who provided us with some of the samples used. For the CTA strains: João Aristeu da Rosa (Universidade Estadual Paulista, Unesp), Faculdade de Ciências Farmacêuticas, Araraquara, São Paulo, Brazil); and for *R. prolixus* strain from Honduras: Claudio Lazzari (Université de Tours, IRBI, France). We would also like to thank Claire Capdevielle-Dulac and Léa Payen for their help in molecular biology experiments. We thank the GenOuest bioinformatic platform for computing resources and technical support. We thank the Genotoul sequencing facility for technical support and particularly Carole Lampietro for her assistance.

## Authors Contribution

JF, MH, and CEA designed the project;, CEA, E F-R, and MH processed the samples; GR and MM carried out the bioinformatics analyses with the help of JF ; GR, MM, MH and JF analyzed the data; CEA and EF-R provided expertise on the biology of hematophagous triatomines; GR, MH and JF wrote the article. All the authors reviewed and edited the manuscript.

## Data and Code Availability

The raw sequencing data generated in this study have been deposited in the NCBI Sequence Read Archive (SRA) under the NCBI BioProject accession [PRJNA1405103](https://www.ncbi.nlm.nih.gov/bioproject/1405103). Raw data and other supplementary datasets are available through the Zenodo community at: https://zenodo.org/communities/kissingbugsomics/. BlobTools analyses, phylogenies, host assemblies, and metabolism analyses are available at: https://doi.org/10.5281/zenodo.18354111, and the *Triatoma brasiliensis* genome assembly is available at: https://doi.org/10.5281/zenodo.19135825. SisGen registration: A5C8D0D.

