## Supplementary Figures and Table for "Metagenomic Analyses Reveal a Constrained Network of Nutritional Symbionts in Kissing Bugs"

**Supplementary Figure 1: Detection of symbionts within the metagenomes of nine Triatominae species:** Contigs represented as circles were binned based on their GC%, read coverage, and taxonomic assignment. The size of contigs is represented through the size of the circles

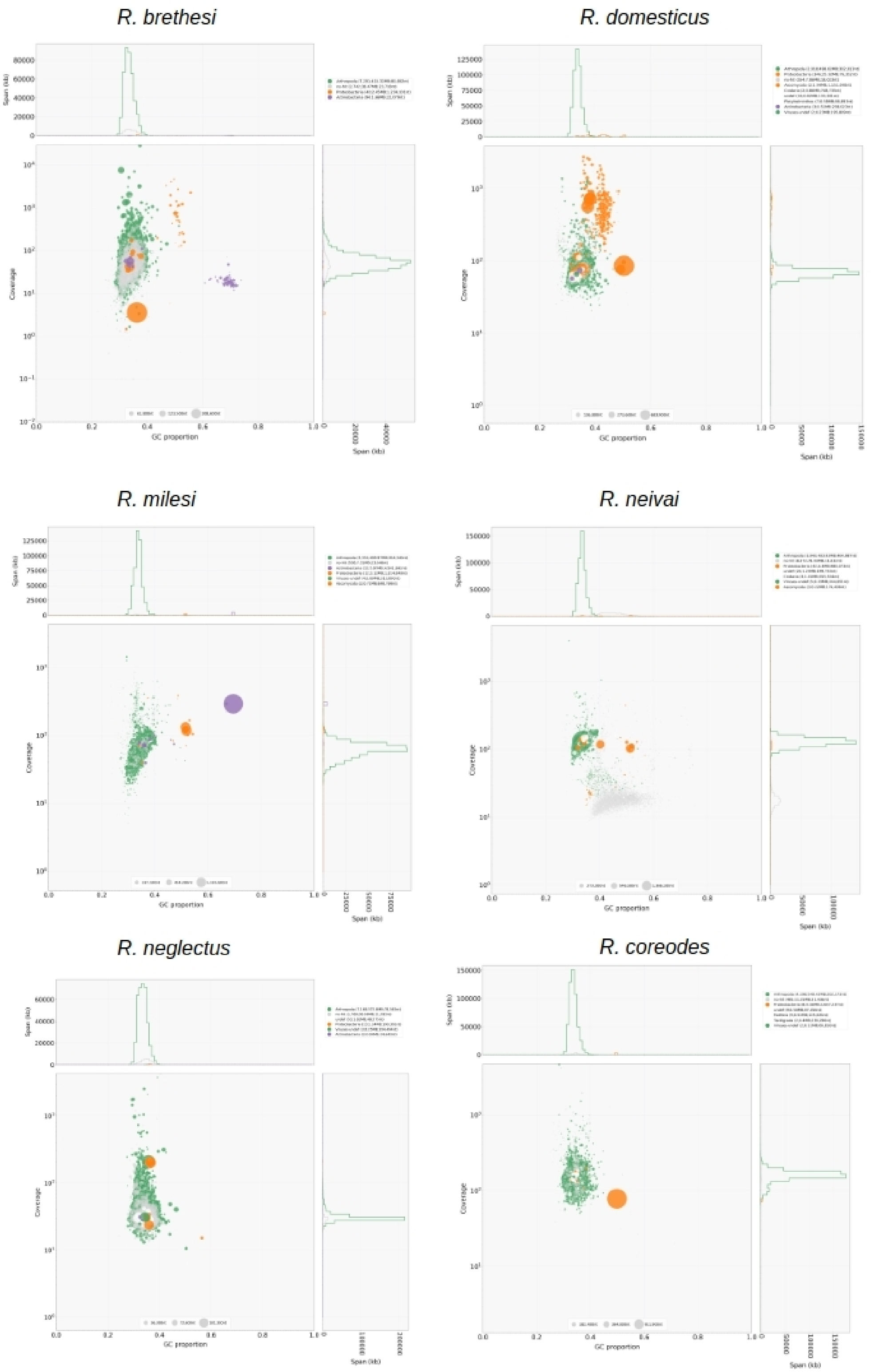

*R. pictipes*

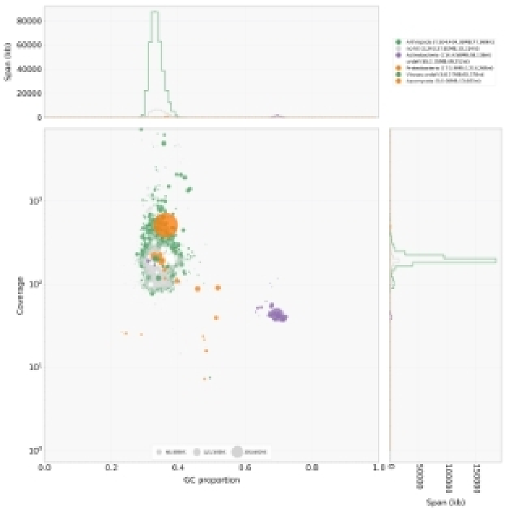

*R. robustus*

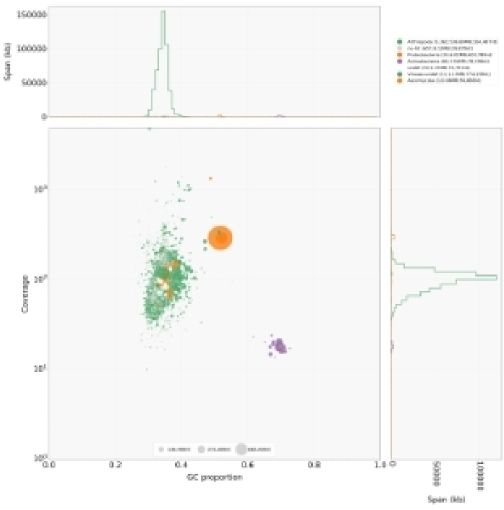

*T. brasiliensis*

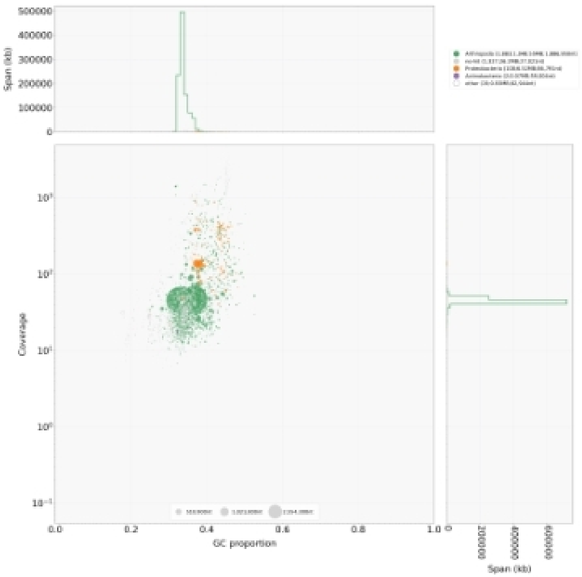

**Supplementary Figure 2: Putative functions of the laterally transferred genes found in the host insect genomes.**

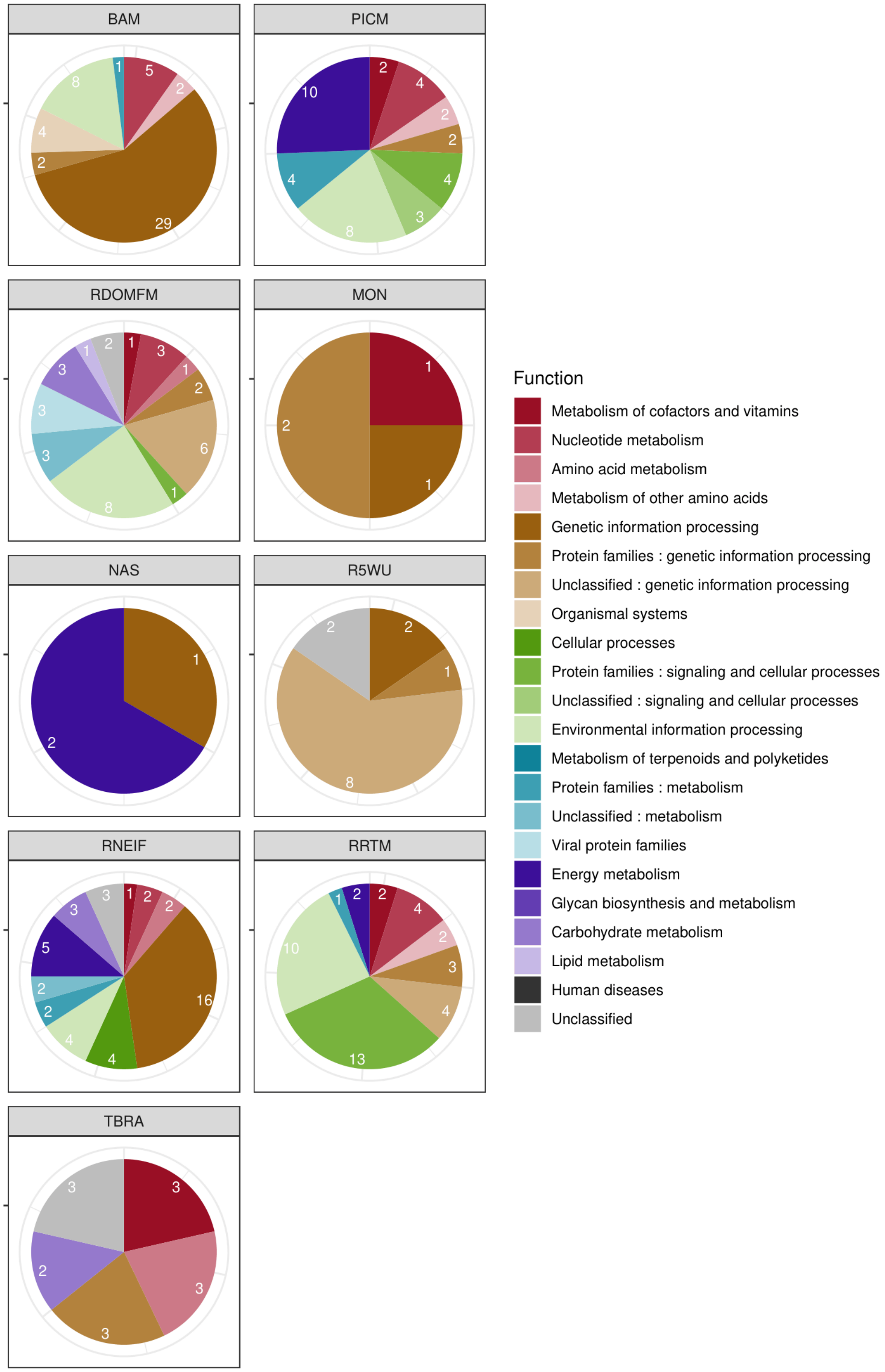

**Supplementary Figure 3 : Whole *Wolbachia* gene phylogeny of Triatominae-associated symbionts and reference genomes.** The tree represents the best maximum-likelihood tree obtained from 44 conserved single-copy orthologs. Ultrafast bootstrap scores (from 1,000 replicates) above 80 are indicated in red below the relevant node. The colored range shows the group formed by Triatominae-associated symbionts.

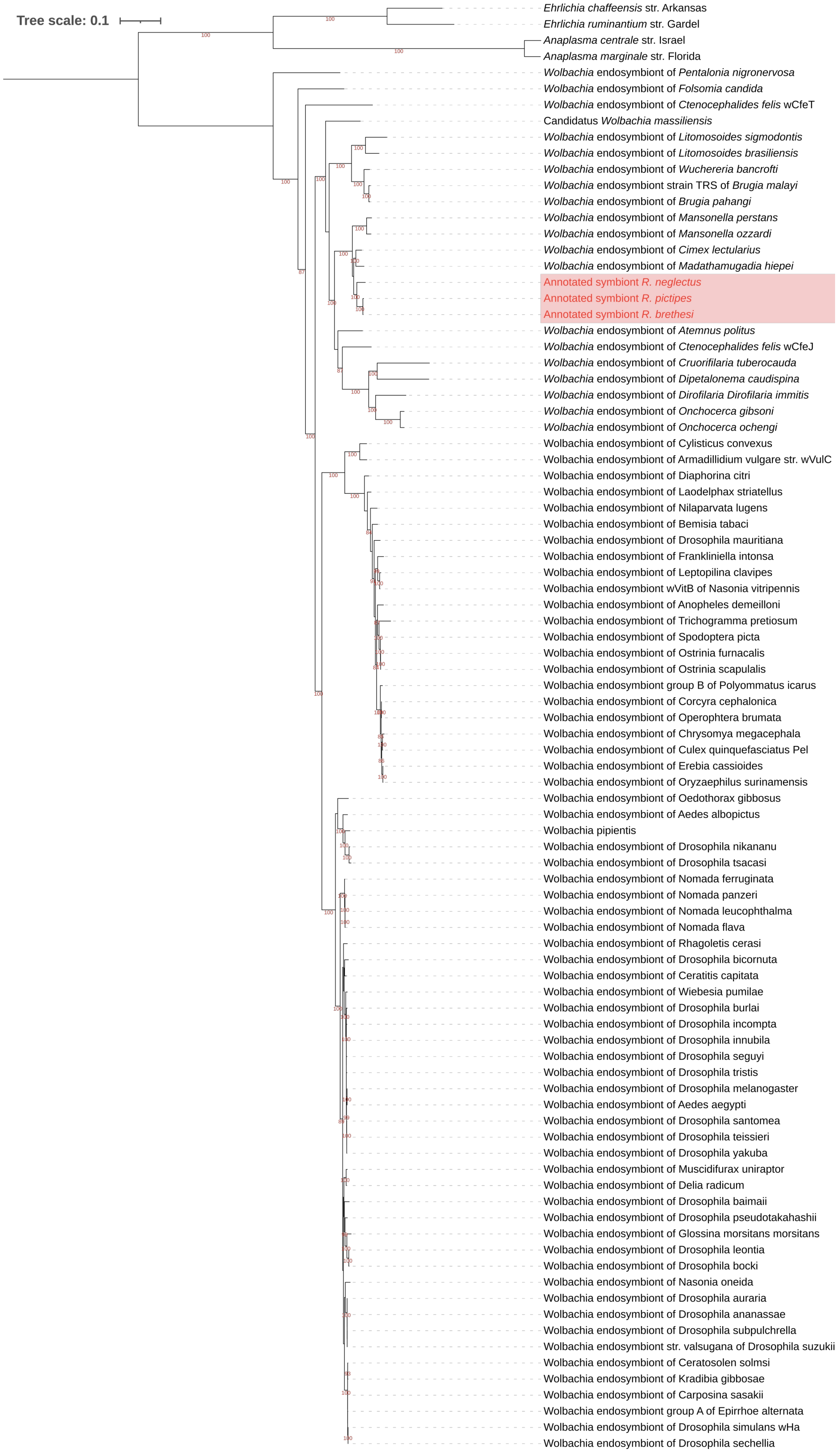

**Supplementary Figure 4: Functional analysis of the different genes found in the two *Arsenophonus* symbionts associated with *R. domesticus***. The upper and middle panel indicate the gene function predictions of the largest and the smaller *Arsenophonus* genome respectively, whereas the lower panel describe the functional predictions of the genes that have undergone gene deletions in the smaller assembly.

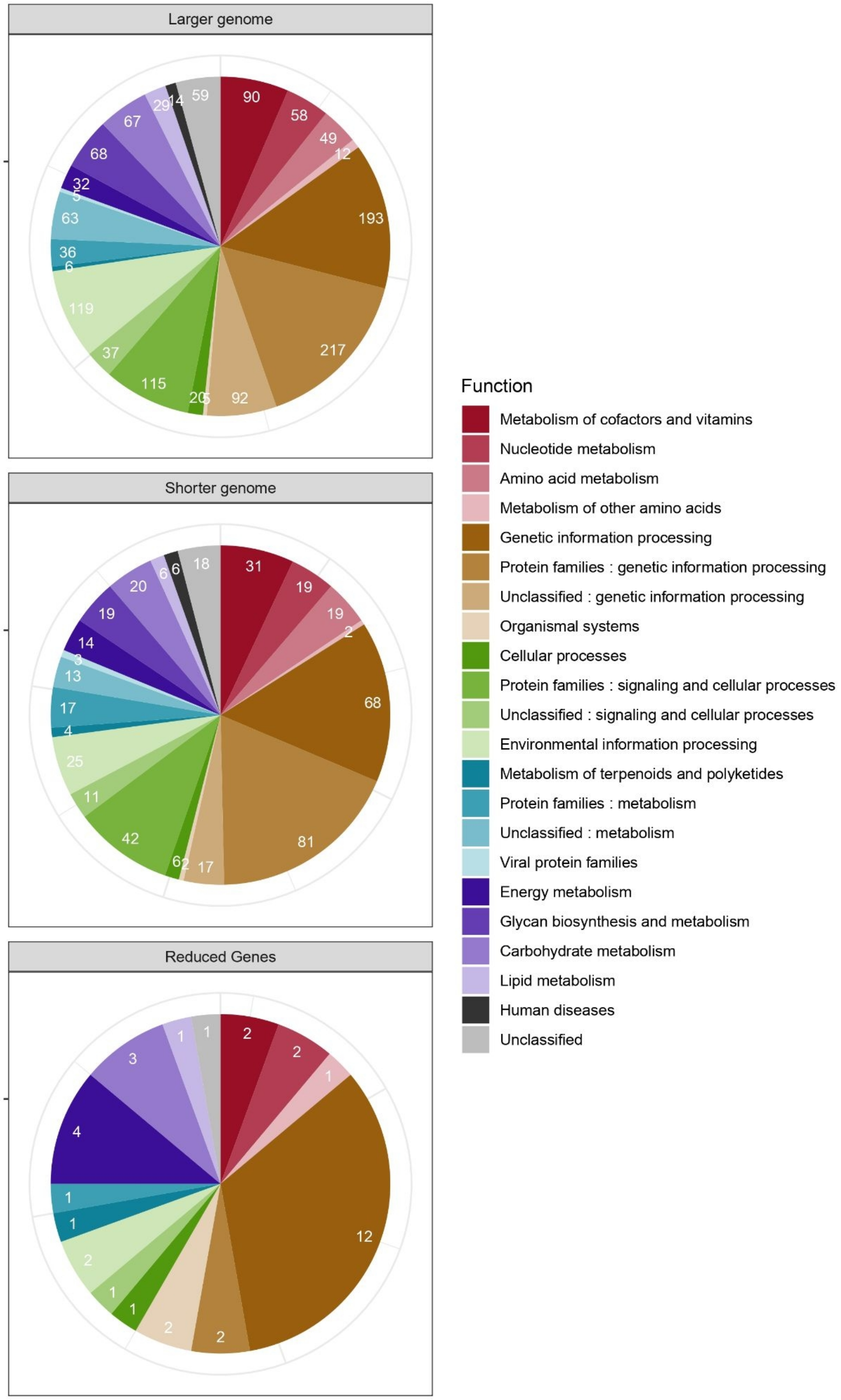

**Supplementary Table 1: List of the 165 reference genomes used for the phylogeny analysis**

| Accession | Assignation | Host/medium | Contigs | Genome size |
|---|---|---|---|---|
| GCF_020268605.1 | *Arsenophonus apicola ArsBeeUS* | *Apis melifera* | 7 | 3,60E+06 |
| GCF_029906405.1 | *Arsenophonus apicola aApi AU* | *Apis melifera* | 6 | 3,30E+06 |
| GCF_903968575.1 | *Arsenophonus endosymbiont of Apis melifera* | *Apis melifera* | 79 | 3,30E+06 |
| GCF_004768525.1 | *Arsenophonus nasoniae FIN* | *Nasonia vitripennis* | 18 | 5,00E+06 |
| GCF_029873515.1 | *Arsenophonus nasoniae aNv CAN* | *Nasonia vitripennis* | 21 | 5,10E+06 |
| GCF_029873555.1 | *Arsenophonus nasoniae aNVv UK* | *Nasonia vitripennis* | 17 | 4,70E+06 |
| GCF_029873535.1 | *Arsenophonus nasoniae aNv CH* | *Nasonia vitripennis* | 17 | 4,70E+06 |
| GCF_029873495.1 | *Arsenophonus nasoniae aPv* | *Nasonia vitripennis* | 19 | 4,30E+06 |
| GCF_029873455.1 | *Arsenophonus nasoniae alh* | *Nasonia vitripennis* | 8 | 3,70E+06 |
| GCF_000429565.1 | *Arsenophonus nasoniae DSM 15247* | *Nasonia vitripennis* | 296 | 3,70E+06 |
| GCF_013460135.1 | *Arsenophonus endosymbiont of Aphis craccivora* | *Aphis craccivora* | 2 | 2,40E+06 |
| GCF_900343025.1 | *Arsenophonus endosymbiont of Aleurodicus floccissimus* | *Aleurodicus floccissimus* | 11 | 3,00E+06 |

| | | | | |
|---|---|---|---|---|
| GCF_001640365.1 | *Candidatus Arsenophonus triatominarum* | *Triatoma infestans* | 94 | 3,90E+06 |
| GCF_000757905.1 | *Candidatus Arsenophonus nilaparvatae str. Hanzhou* | *Nilaparvata lugens* | 20 | 3,00E+06 |
| GCF_902713415.1 | *Arsenophonus endosymbiont of Bernisia tabaci Q2* | *Bernisia tabaci* | 328 | 1,90E+06 |
| GCF_029873475.1 | *Arsenophonus sp. aPb* | *Lysandra bellargus* | 5 | 3,50E+06 |
| GCF_002287155.1 | *Arsenophonus sp. ENCA* | *Entylia carinata* | 818 | 3,20E+06 |
| GCA_031432035.1 | *Arsenophonus sp. K10D* | *Pseudolynchia canariensis* | 354 | 1,20E+06 |
| GCF_031432075.1 | *Arsenophonus sp. 20K* | *Crataerina pallida* | 333 | 3,10E+06 |
| GCF_031432055.1 | *Arsenophonus sp. 24K* | *Crataerina pallida* | 167 | 2,80E+06 |
| GCF_031432015.1 | *Arsenophonus sp. K28* | *Hippobosca equina* | 16 | 3,20E+06 |
| GCF_000069965.1 | *Proteus mirabilis* | *Homo sapiens* | 2 | 4,10E+06 |
| GCF_006094455.1 | *Morganella morganii* | *Homo sapiens* | 1 | 3,90E+06 |
| GCF_900478115.1 | *Rhodococcus coprophilus* | *Soil* | 1 | 4,58E+06 |
| GCF_017068395.1 | *Rhodococcus pseudokoreensis* | *Soil* | 6 | 9,87E+06 |
| GCF_003130705.1 | *Rhodococcus oxybenzonivorans* | *Soil* | 4 | 8,01E+06 |
| GCF_026153295.1 | *Rhodococcus antarcticus* | *Soil* | 4 | 3,94E+06 |
| GCF_000696675.2 | *Rhodococcus erythropolis R138* | *Soil* | 3 | 6,81E+06 |
| GCF_015099595.1 | *Rhodococcus qingshengii* | *Soil* | 2 | 6,73E+06 |

| | | | | |
|---|---|---|---|---|
| GCF_000511305.1 | *Rhodococcus pyridinivorans SB3094* | *Soil* | 3 | 5,59E+06 |
| GCF_020542785.1 | *Rhodococcus opacus PD630* | *Soil* | 4 | 9,17E+06 |
| GCF_016804345.1 | *Rhodococcus ruber* | *Soil* | 3 | 5,70E+06 |
| GCF_025722975.1 | *Rhodococcus aetherivorans* | *Soil* | 3 | 6,56E+06 |
| GCF_001620305.1 | *Rhodococcus fascians D188* | *Chrysanthemum morifolium* | 3 | 5,50E+06 |
| GCF_019334125.1 | *Rhodococcus globerulus* | *Soil* | 1 | 6,74E+06 |
| GCF_017068375.1 | *Rhodococcus koreensis* | *Soil* | 5 | 1,09E+07 |
| GCF_003004765.2 | *Rhodococcus rhodochrous* | *Soil* | 1 | 5,72E+06 |
| GCF_001895025.1 | *Rhodococcus zopfii NBRC 100606 = JCM 9919* | *Bioréacteur* | 146 | 6,30E+06 |
| GCF_008011915.1 | *Rhodococcus rhodnii* | *Rhodnius prolixus* | 4 | 4,49E+06 |
| GCF_001894765.1 | *Rhodococcus corynebacterioides NBRC 14404* | *Air* | 14 | 3,98E+06 |
| GCF_001894865.1 | *Rhodococcus maanshanensis NBRC 100610* | *Soil* | 97 | 5,67E+06 |
| GCF_001895005.1 | *Rhodococcus yunnanensis NBRC 103083* | *Soil* | 68 | 6,37E+06 |
| GCF_001894885.1 | *Rhodococcus marinonascens NBRC 14363* | *Soil* | 156 | 4,92E+06 |
| GCF_900455725.1 | *Rhodococcus gordoniae* | *Homo sapiens* | 3 | 4,87E+06 |
| GCF_014635345.1 | *Rhodococcus trifolii* | *Trifolium repens* | 15 | 5,29E+06 |
| GCF_004011835.1 | *Rhodococcus spongiicola* | *Marine sponge* | 29 | 3,97E+06 |

| | | | | |
|---|---|---|---|---|
| GCF_004011825.1 | *Rhodococcus xishaensis* | *Marine sponge* | 22 | 3,71E+06 |
| GCF_005049235.1 | *Rhodococcus oryzae* | *Soil* | 31 | 5,37E+06 |
| GCF_006704125.1 | *Rhodococcus spelaei* | *Soil* | 18 | 4,81E+06 |
| GCF_001646785.1 | *Rhodococcus phenolicus* | *Soil* | 232 | 6,28E+06 |
| GCF_020515525.1 | *Rhodococcus yananensis* | *Fermentation bed* | 178 | 4,25E+06 |
| GCF_030509825.1 | *Rhodococcus electrodiphilus* | *Coral* | 225 | 5,49E+06 |
| GCA_003096095.1 | *Dietzia psychralcaliphila* | *Water* | 2 | 3,90E+06 |
| GCA_027270575.1 | *Dietzia maris* | *Soil* | 84 | 3,62E+06 |
| GCA_000092225.1 | *Tsukamurella paurometabola DSM 20162* | *Homo sapiens* | 2 | 4,48E+06 |
| GCA_900460155.1 | *Tsukamurella pulmonis* | *Homo sapiens* | 2 | 4,77E+06 |
| GCA_019797845.1 | *Symbiopectobacterium purcellii* | *Empoasca decipiens* | 1 | 4,94E+06 |
| GCA_018777385.1 | *Candidatus Symbiopectobacterium endolongispinus* | *Pseudococcus longispinus* | 10 | 4,49E+06 |
| GCA_025962675.1 | *Candidatus Symbiopectobacterium sp. NZEC127* | *Potato tubers* | 116 | 5,29E+06 |
| GCA_025962655.1 | *Candidatus Symbiopectobacterium sp. NZEC151* | *Potato tubers* | 36 | 5,16E+06 |
| GCA_015762685.1 | *Candidatus Symbiopectobacterium sp. 'North America'* | *Howardula aoronymphium* | 125 | 4,56E+06 |

| | | | | |
|---|---|---|---|---|
| GCA_025962695.1 | *Candidatus Symbiopectobacterium sp. NZEC135* | *Potato tubers* | 1588 | 6,70E+06 |
| GCA_015762695.1 | *Candidatus Symbiopectobacterium sp. PLON1* | *Pseudococcus longispinus* | 83 | 4,41E+06 |
| GCA_015762725.1 | *Candidatus Symbiopectobacterium sp. Dall1.0* | *Diachasma alloeum* | 500 | 3,64E+06 |
| GCF_020054415.1 | *Candidatus Symbiopectobacterium sp.* | *Rhodnius prolixus* | 743 | 3,29E+06 |
| GCA_012689415.1 | *Candidatus Symbiopectobacterium sp. Chty_BC* | *Chilacis typhae* | 632 | 1,50E+06 |
| GCF_001742185.1 | *Pectobacterium_wasabiae* | *Eutrema japonicum* | 1 | 5,04E+06 |
| GCF_013488025.1 | *Pectobacterium_carotovorum* | *Potato* | 1 | 4,89E+06 |
| GCF_000740965.1 | *Pectobacterium_atrosepticum* | *Solanum tuberosum* | 2 | 5,02E+06 |
| GCF_000147055.1 | *Dickeya_dadantii_3937* | *Potato* | 1 | 4,92E+06 |
| GCF_002887555.1 | *Dickeya_zeae* | *Banana* | 1 | 4,74E+06 |
| GCF_016584425.1 | *Wolbachia endosymbiont of Drosophila melanogaster* | *Drosophila melanogaster* | 1 | 1,27E+06 |
| GCF_000008385.1 | *Wolbachia endosymbiont strain TRS of Brugia malayi* | *Brugia malayi* | 1 | 1,08E+06 |
| GCF_029856955.1 | *Wolbachia endosymbiont of Frankliniella intonsa* | *Frankliniella intonsa* | 1 | 1,46E+06 |
| GCF_008033215.1 | *Wolbachia endosymbiont of Drosophila ananassae* | *Drosophila ananassae* | 1 | 1,40E+06 |

| | | | | |
|---|---|---|---|---|
| GCF_000376605.1 | *Wolbachia endosymbiont of Drosophila simulans* | *Drosophila simulans wHa* | 1 | 1,30E+06 |
| GCF_013365435.1 | *Wolbachia endosymbiont of Litomosoides sigmodontis* | *Litomosoides sigmodontis* | 1 | 1,05E+06 |
| GCF_000306885.1 | *Wolbachia endosymbiont of Onchocerca ochengi* | *Onchocerca ochengi* | 1 | 9,58E+05 |
| GCF_001931755.2 | *Wolbachia endosymbiont of Folsomia candida* | *Folsomia candida* | 1 | 1,80E+06 |
| GCF_025617515.1 | *Wolbachia endosymbiont of Oryzaephilus surinamensis* | *Oryzaephilus surinamensis* | 1 | 1,73E+06 |
| GCF_021609905.1 | *Wolbachia endosymbiont of Delia radicum* | *Delia radicum* | 1 | 1,59E+06 |
| GCF_947250745.1 | *Wolbachia endosymbiont (group B) of Polyommatus icarus* | *Polyommatus icarus* | 1 | 1,57E+06 |
| GCF_936270145.1 | *Wolbachia endosymbiont of Oedothorax gibbosus* | *Oedothorax gibbosus* | 1 | 1,55E+06 |
| GCF_012277295.1 | *Wolbachia endosymbiont of Ctenocephalides felis wCfeT* | *Ctenocephalides felis* | 1 | 1,50E+06 |
| GCF_006542295.1 | *Wolbachia endosymbiont of Carposina sasakii* | *Carposina sasakii* | 1 | 1,45E+06 |
| GCF_018467135.1 | *Wolbachia endosymbiont of Drosophila santomea* | *Drosophila santomea* | 1 | 1,41E+06 |

| | | | | |
|---|---|---|---|---|
| GCF_018467115.1 | *Wolbachia endosymbiont of Drosophila yakuba* | *Drosophila yakuba* | 1 | 1,39E+06 |
| GCF_008245065.1 | *Wolbachia endosymbiont of Chrysomya megacephala* | *Chrysomya megacephala* | 1 | 1,38E+06 |
| GCF_020995475.1 | *Wolbachia endosymbiont of Corcyra cephalonica* | *Corcyra cephalonica* | 3 | 1,38E+06 |
| GCF_018141665.1 | *Wolbachia endosymbiont of Spodoptera picta* | *Spodoptera picta* | 1 | 1,34E+06 |
| GCF_023559125.1 | *Wolbachia endosymbiont of Ostrinia furnacalis* | *Ostrinia furnacalis* | 1 | 1,32E+06 |
| GCF_023559145.1 | *Wolbachia endosymbiont of Ostrinia scapulalis* | *Ostrinia scapulalis* | 1 | 1,32E+06 |
| GCF_026015925.1 | *Wolbachia endosymbiont of Drosophila pseudotakahashii* | *Drosophila pseudotakahashii* | 1 | 1,31E+06 |
| GCF_021378375.1 | *Wolbachia endosymbiont of Drosophila innubila* | *Drosophila innubila* | 1 | 1,29E+06 |
| GCF_017869285.1 | *Wolbachia endosymbiont of Wiebesia pumilae* | *Wiebesia pumilae* | 1 | 1,28E+06 |
| GCF_004795975.1 | *Wolbachia endosymbiont of Drosophila mauritiana* | *Drosophila mauritiana* | 1 | 1,27E+06 |
| GCF_017896245.1 | *Wolbachia endosymbiont of Aedes aegypti* | *Aedes aegypti* | 1 | 1,27E+06 |
| GCF_000829315.1 | *Wolbachia endosymbiont of Cimex lectularius* | *Cimex lectularius* | 1 | 1,25E+06 |

| | | | | |
|---|---|---|---|---|
| GCF_018491735.2 | *Wolbachia endosymbiont of Anopheles demeilloni* | *Anopheles demeilloni* | 1 | 1,23E+06 |
| GCF_012277315.1 | *Wolbachia endosymbiont of Ctenocephalides felis wCfeJ* | *Ctenocephalides felis* | 1 | 1,20E+06 |
| GCF_947251475.1 | *Wolbachia endosymbiont (group A) of Epirrhoe alternata* | *Epirrhoe alternata* | 1 | 1,20E+06 |
| GCF_024804185.1 | *Wolbachia endosymbiont of Aedes albopictus* | *Aedes albopictus* | 3 | 1,23E+06 |
| GCF_012030695.1 | *Wolbachia endosymbiont of Brugia pahangi* | *Brugia pahangi* | 1 | 1,07E+06 |
| GCF_013365455.1 | *Wolbachia endosymbiont of Dirofilaria (Dirofilaria) immitis* | *Dirofilaria immitis* | 1 | 9,20E+05 |
| GCF_013365475.1 | *Wolbachia endosymbiont of Cruorifilaria tuberocauda* | *Cruorifilaria tuberocauda* | 1 | 8,64E+05 |
| GCF_013365495.1 | *Wolbachia endosymbiont of Dipetalonema caudispina* | *Dipetalonema caudispina* | 1 | 8,63E+05 |
| GCF_014107475.1 | *Wolbachia pipientis* | *Drosophila sturtevanti* | 1 | 1,19E+06 |
| GCF_013096725.2 | *Wolbachia endosymbiont of Diaphorina citri* | *Diaphorina citri* | 1 | 1,52E+06 |
| GCF_000073005.1 | *Wolbachia endosymbiont of Culex quinquefasciatus Pel* | *Culex quinquefasciatus* | 1 | 1,48E+06 |

| | | | | |
|---|---|---|---|---|
| GCF_017869155.1 | *Wolbachia endosymbiont of Ceratosolen solmsi* | *Ceratosolen solmsi* | 1 | 1,21E+06 |
| GCF_003999585.1 | *Wolbachia endosymbiont of Bemisia tabaci* | *Bemisia tabaci* | 1 | 1,31E+06 |
| GCF_014771645.1 | *Candidatus Wolbachia massiliensis* | *Cimex hemipterus* | 1 | 1,29E+06 |
| GCF_001758565.1 | *Wolbachia endosymbiont of Drosophila incompta* | *Drosophila incompta* | 1 | 1,27E+06 |
| GCF_001439985.1 | *Wolbachia endosymbiont of Trichogramma pretiosum* | *Trichogramma pretiosum* | 1 | 1,13E+06 |
| GCF_007115015.1 | *Wolbachia endosymbiont of Laodelphax striatellus* | *Laodelphax striatellus* | 2 | 1,79E+06 |
| GCF_002204235.2 | *Wolbachia endosymbiont of Wuchereria bancrofti* | *Wuchereria bancrofti* | 100 | 1,06E+06 |
| GCF_000174095.1 | *Wolbachia endosymbiont of Muscidifurax uniraptor* | *Muscidifurax uniraptor* | 256 | 8,68E+05 |
| GCF_000204545.1 | *Wolbachia endosymbiont wVitB of Nasonia vitripennis* | *Nasonia vitripennis* | 426 | 1,11E+06 |
| GCF_000333795.1 | *Wolbachia endosymbiont str. valsugana of Drosophila suzukii* | *Drosophila suzukii* | 110 | 1,42E+06 |
| GCF_018555315.1 | *Wolbachia endosymbiont of Erebia cassioides* | *Erebia cassioides* | 2 | 1,42E+06 |

| | | | | |
|---|---|---|---|---|
| GCF_016031645.1 | *Wolbachia endosymbiont of Kradibia gibbosae* | *Kradibia gibbosae* | 3 | 1,45E+06 |
| GCF_001027565.1 | *Wolbachia endosymbiont of Armadillidium vulgare str. wVulC* | *Armadillidium vulgare* | 10 | 1,66E+06 |
| GCF_007115045.1 | *Wolbachia endosymbiont of Nilaparvata lugens* | *Nilaparvata lugens* | 2 | 1,54E+06 |
| GCF_018454475.1 | *Wolbachia endosymbiont of Rhagoletis cerasi* | *Rhagoletis cerasi* | 16 | 1,26E+06 |
| GCF_029169405.1 | *Wolbachia endosymbiont of Onchocerca gibsoni* | *Onchocerca gibsoni* | 1 | 9,98E+05 |
| GCF_013366805.1 | *Wolbachia endosymbiont of Litomosoides brasiliensis* | *Litomosoides brasiliensis* | 41 | 1,05E+06 |
| GCF_006334525.1 | *Wolbachia endosymbiont of Leptopilina clavipes* | *Leptopilina clavipes* | 46 | 1,15E+06 |
| GCF_009012935.1 | *Wolbachia endosymbiont of Nasonia oneida* | *Nasonia oneida* | 47 | 1,29E+06 |
| GCF_018454455.1 | *Wolbachia endosymbiont of Ceratitis capitata* | *Ceratitis capitata* | 65 | 1,24E+06 |
| GCF_002300525.1 | *Wolbachia endosymbiont of Drosophila subpulchrella* | *Drosophila subpulchrella* | 106 | 1,42E+06 |
| GCF_028982115.1 | *Wolbachia endosymbiont of Drosophila tsacasi* | *Drosophila tsacasi* | 59 | 1,13E+06 |

| | | | | |
|---|---|---|---|---|
| GCF_028981865.1 | *Wolbachia endosymbiont of Drosophila nikananu* | *Drosophila nikananu* | 83 | 1,26E+06 |
| GCF_014354315.1 | *Wolbachia endosymbiont of Drosophila sechellia* | *Drosophila sechellia* | 83 | 1,29E+06 |
| GCF_003344345.1 | *Wolbachia endosymbiont of Cylisticus convexus* | *Cylisticus convexus* | 237 | 2,11E+06 |
| GCF_028981745.1 | *Wolbachia endosymbiont of Drosophila tristis* | *Drosophila tristis* | 114 | 1,26E+06 |
| GCF_001675695.1 | *Wolbachia endosymbiont of Nomada flava* | *Nomada flava* | 167 | 1,33E+06 |
| GCF_001675715.1 | *Wolbachia endosymbiont of Nomada leucophthalma* | *Nomada leucophthalma* | 182 | 1,37E+06 |
| GCF_001675775.1 | *Wolbachia endosymbiont of Nomada panzeri* | *Nomada panzeri* | 191 | 1,34E+06 |
| GCF_017916175.1 | *Wolbachia endosymbiont of Drosophila auraria* | *Drosophila auraria* | 116 | 1,32E+06 |
| GCF_028981925.1 | *Wolbachia endosymbiont of Drosophila burlai* | *Drosophila burlai* | 128 | 1,25E+06 |
| GCF_014534705.1 | *Wolbachia endosymbiont of Pentalonia nigronervosa* | *Pentalonia nigronervosa* | 182 | 1,46E+06 |
| GCF_005862135.1 | *Wolbachia endosymbiont of Drosophila teissieri* | *Drosophila teissieri* | 122 | 1,30E+06 |
| GCF_020278625.1 | *Wolbachia endosymbiont of Mansonella ozzardi* | *Mansonella ozzardi* | 93 | 1,07E+06 |

| | | | | |
|---|---|---|---|---|
| GCF_001675785.1 | *Wolbachia endosymbiont of Nomada ferruginata* | *Nomada ferruginata* | 231 | 1,34E+06 |
| GCF_028982105.1 | *Wolbachia endosymbiont of Drosophila bicornuta* | *Drosophila bicornuta* | 140 | 1,19E+06 |
| GCF_001266585.1 | *Wolbachia endosymbiont of Operophtera brumata* | *Operophtera brumata* | 120 | 1,12E+06 |
| GCF_013309895.1 | *Wolbachia endosymbiont of Atemnus politus* | *Atemnus politus* | 200 | 1,40E+06 |
| GCF_020278605.1 | *Wolbachia endosymbiont of Mansonella perstans* | *Mansonella perstans* | 170 | 1,06E+06 |
| GCF_028981765.1 | *Wolbachia endosymbiont of Drosophila seguyi* | *Drosophila seguyi* | 201 | 1,17E+06 |
| GCF_028982185.1 | *Wolbachia endosymbiont of Drosophila baimaii* | *Drosophila baimaii* | 209 | 1,14E+06 |
| GCF_028981785.1 | *Wolbachia endosymbiont of Drosophila leontia* | *Drosophila leontia* | 186 | 1,10E+06 |
| GCF_013366855.1 | *Wolbachia endosymbiont of Madathamugadia hiepei* | *Madathamugadia hiepei* | 208 | 1,03E+06 |
| GCF_028982005.1 | *Wolbachia endosymbiont of Drosophila bocki* | *Drosophila bocki* | 227 | 1,09E+06 |
| GCF_000689175.1 | *Wolbachia endosymbiont of Glossina morsitans morsitans* | *Glossina morsitans* | 201 | 1,02E+06 |
| GCF_000024505.1 | *Anaplasma centrale str. Israel* | *Cattle* | 1 | 1,21E+06 |

| | | | | |
|---|---|---|---|---|
| GCF_000020305.1 | *Anaplasma marginale str. Florida* | *Cattle* | 1 | 1,20E+06 |
| GCF_000013145.1 | *Ehrlichia chaffeensis str. Arkansas* | *Homo sapiens* | 1 | 1,18E+06 |
| GCF_000050405.1 | *Ehrlichia ruminantium str. Gardel* | *Homo sapiens* | 1 | 1,50E+06 |

**Supplementary table 2 : List of all reference genes used for the metabolic pathway search.** Genes involved in multiple pathways are only indicated once and referenced with both pathways.

| Gene accession | Organism | Pathway |
|---|---|---|
| LY180_00870 | *E. coli* | Biotin |
| LY180_04075 | | |
| LY180_04080 | | |
| LY180_04085 | | |
| LY180_04090 | | |
| LY180_04095 | | |
| LY180_05670 | | |
| LY180_05680 | | |
| LY180_06560 | | |
| LY180_08295 | | |
| LY180_12035 | | |
| LY180_15420 | | |
| LY180_17500 | | |
| LY180_18240 | | |
| LY180_20845 | | |
| ArsFIN_06280 | *A. nasoniae* | |
| ArsFIN_08000 | | |
| ArsFIN_15560 | | |
| ArsFIN_15570 | | |
| ArsFIN_15580 | | |
| ArsFIN_15590 | | |
| ArsFIN_15600 | | |
| ArsFIN_21230 | | |

| | |
|---|---|
| ArsFIN_24940 | |
| ArsFIN_24960 | |
| ArsFIN_26790 | |
| ArsFIN_41270 | |
| KYT97_01805 | *R. globerulus* |
| KYT97_03030 | |
| KYT97_06065 | |
| KYT97_06255 | |
| KYT97_06260 | |
| KYT97_06265 | |
| KYT97_07900 | |
| KYT97_12585 | |
| KYT97_13390 | |
| KYT97_13600 | |
| KYT97_14080 | |
| KYT97_16100 | |
| KYT97_16560 | |
| KYT97_17245 | |
| KYT97_17460 | |
| KYT97_18880 | |
| KYT97_21540 | |
| KYT97_23580 | |
| K6K13_07705 | *S. purcellii* |
| K6K13_07710 | |
| K6K13_07715 | |
| K6K13_07720 | |

| | | |
|---|---|---|
| K6K13_07725 | | |
| K6K13_09115 | | |
| K6K13_12775 | | |
| K6K13_12785 | | |
| K6K13_14185 | | |
| K6K13_19035 | | |
| K6K13_21750 | | |
| K6K13_22600 | | |
| K6K13_22840 | | |
| WCLE_001810 | *W. C. lectularius* | |
| WCLE_004670 | | |
| WCLE_008500 | | |
| WCLE_008510 | | |
| WCLE_008520 | | |
| WCLE_008540 | | |
| WCLE_008550 | | |
| WCLE_008830 | | |
| WCLE_009890 | | |
| WCLE_010330 | | |
| LY180_06495 | *E. coli* | Riboflavin/ Folate |
| ArsFIN_19690 | *A. nasoniae* | |
| K6K13_09045 | *S. purcellii* | |
| LY180_11000 | *E. coli* | Riboflavin/ Thiamine |
| K6K13_15030 | *S. purcellii* | |
| LY180_00125 | *E. coli* | Riboflavin |
| LY180_02400 | | |

| | |
|---|---|
| LY180_02405 | |
| LY180_04445 | |
| LY180_04925 | |
| LY180_05140 | |
| LY180_05875 | |
| LY180_08665 | |
| LY180_11975 | |
| LY180_13915 | |
| LY180_15675 | |
| LY180_15710 | |
| LY180_19740 | |
| LY180_19910 | |
| LY180_21300 | |
| LY180_22795 | |
| ArsFIN_02400 | *A. nasoniae* |
| ArsFIN_05530 | |
| ArsFIN_05570 | |
| ArsFIN_17230 | |
| ArsFIN_24490 | |
| ArsFIN_26670 | |
| ArsFIN_34960 | |
| ArsFIN_34970 | |
| ArsFIN_35210 | |
| ArsFIN_37690 | |
| K6K13_01480 | *S. purcellii* |
| K6K13_04775 | |

| | | |
|---|---|---|
| K6K13_08955 | | |
| K6K13_11310 | | |
| K6K13_14120 | | |
| K6K13_18235 | | |
| K6K13_18450 | | |
| K6K13_18455 | | |
| K6K13_19845 | | |
| K6K13_22075 | | |
| LY180_00020 | *E. coli* | B6 |
| LY180_00265 | | |
| LY180_04765 | | |
| LY180_07325 | | |
| LY180_08535 | | |
| LY180_08545 | | |
| LY180_12020 | | |
| LY180_12445 | | |
| LY180_13155 | | |
| LY180_15065 | | |
| ArsFIN_05680 | *A. nasoniae* | |
| ArsFIN_12990 | | |
| ArsFIN_21380 | | |
| ArsFIN_21400 | | |
| ArsFIN_26760 | | |
| ArsFIN_29630 | | |
| ArsFIN_35490 | | |
| ArsFIN_36460 | | |

| | | |
|---|---|---|
| K6K13_05455 | *S. purcellii* | |
| K6K13_09020 | | |
| K6K13_09030 | | |
| K6K13_12310 | | |
| K6K13_14165 | | |
| K6K13_15510 | | |
| K6K13_18150 | | |
| K6K13_18300 | | |
| LY180_02240 | *E. coli* | Thiamine/ Folate |
| K6K13_21735 | *S. purcellii* | |
| LY180_02415 | *E. coli* | Thiamine |
| LY180_02430 | | |
| LY180_02445 | | |
| LY180_02710 | | |
| LY180_05735 | | |
| LY180_10995 | | |
| LY180_12970 | | |
| LY180_20920 | | |
| LY180_20925 | | |
| LY180_20935 | | |
| LY180_20940 | | |
| LY180_20945 | | |
| LY180_21865 | | |
| ArsFIN_04080 | *A. nasoniae* | |
| ArsFIN_16310 | | |
| ArsFIN_24860 | | |

| | | |
|---|---|---|
| ArsFIN_29420 | | |
| ArsFIN_34500 | | |
| ArsFIN_34890 | | |
| ArsFIN_34920 | | |
| ArsFIN_34940 | | |
| ArsFIN_41410 | | |
| ArsFIN_41420 | | |
| ArsFIN_41430 | | |
| ArsFIN_41450 | | |
| ArsFIN_41460 | | |
| K6K13_05040 | | |
| K6K13_05730 | | |
| K6K13_12725 | | |
| K6K13_15025 | | |
| K6K13_15315 | | |
| K6K13_18415 | | |
| K6K13_18430 | *S. purcellii* | |
| K6K13_18440 | | |
| K6K13_22465 | | |
| K6K13_22470 | | |
| K6K13_22475 | | |
| K6K13_22485 | | |
| K6K13_22490 | | |
| LY180_14130 | *E. coli* | Nictotinamide/ Pantothenate |
| KYT97_28740 | *R. globerulus* | |

| | | |
|---|---|---|
| LY180_00525 | *E. coli* | Nictotinamide |
| LY180_02740 | | |
| LY180_03450 | | |
| LY180_03605 | | |
| LY180_03955 | | |
| LY180_04895 | | |
| LY180_05805 | | |
| LY180_08355 | | |
| LY180_08360 | | |
| LY180_09055 | | |
| LY180_09205 | | |
| LY180_13205 | | |
| LY180_13400 | | |
| LY180_13480 | | |
| LY180_13680 | | |
| LY180_13940 | | |
| LY180_20790 | | |
| LY180_20955 | | |
| LY180_22430 | | |
| LY180_23010 | | |
| LY180_23040 | | |
| ArsFIN_08520 | *A. nasoniae* | |
| ArsFIN_09820 | | |
| ArsFIN_10370 | | |
| ArsFIN_12400 | | |
| ArsFIN_13910 | | |

| | |
|---|---|
| ArsFIN_18900 | |
| ArsFIN_18910 | |
| ArsFIN_20060 | |
| ArsFIN_24590 | |
| ArsFIN_29520 | |
| ArsFIN_29730 | |
| ArsFIN_29780 | |
| ArsFIN_31940 | |
| ArsFIN_34430 | |
| ArsFIN_35610 | |
| ArsFIN_35650 | |
| ArsFIN_35940 | |
| ArsFIN_36580 | |
| ArsFIN_37420 | |
| ArsFIN_41400 | |
| KYT97_01250 | *R. globerulus* |
| KYT97_05025 | |
| KYT97_06020 | |
| KYT97_06025 | |
| KYT97_06030 | |
| KYT97_07150 | |
| KYT97_07460 | |
| KYT97_08165 | |
| KYT97_11330 | |
| KYT97_11430 | |
| KYT97_16185 | |

| | | |
|---|---|---|
| KYT97_16310 | | |
| KYT97_20025 | | |
| KYT97_21035 | | |
| KYT97_22375 | | |
| KYT97_23405 | | |
| KYT97_23830 | | |
| KYT97_25975 | | |
| KYT97_26605 | | |
| KYT97_26610 | | |
| KYT97_27090 | | |
| KYT97_27095 | | |
| KYT97_27100 | | |
| KYT97_30615 | | |
| LY180_00380 | *E. coli* | Pantothenate |
| LY180_00385 | | |
| LY180_00500 | | |
| LY180_00635 | | |
| LY180_00645 | | |
| LY180_00650 | | |
| LY180_02345 | | |
| LY180_02455 | | |
| LY180_11220 | | |
| LY180_11225 | | |
| LY180_13150 | | |
| LY180_14785 | | |
| LY180_17825 | | |

| | |
|---|---|
| LY180_18715 | |
| LY180_18740 | |
| LY180_19015 | |
| LY180_19020 | |
| LY180_19515 | |
| LY180_19520 | |
| LY180_19525 | |
| LY180_19530 | |
| LY180_19545 | |
| LY180_20850 | |
| KYT97_00085 | *R. globerulus* |
| KYT97_03170 | |
| KYT97_04405 | |
| KYT97_05260 | |
| KYT97_05355 | |
| KYT97_05675 | |
| KYT97_05735 | |
| KYT97_06135 | |
| KYT97_06140 | |
| KYT97_06755 | |
| KYT97_13095 | |
| KYT97_13475 | |
| KYT97_14695 | |
| KYT97_15385 | |
| KYT97_15430 | |
| KYT97_15790 | |

| | |
|---|---|
| KYT97_19585 | |
| KYT97_22195 | |
| KYT97_22545 | |
| KYT97_22550 | |
| KYT97_22555 | |
| KYT97_23215 | |
| KYT97_23610 | |
| KYT97_23780 | |
| KYT97_25220 | |
| KYT97_25225 | |
| KYT97_25230 | |
| KYT97_25240 | |
| KYT97_25345 | |
| KYT97_26540 | |
| KYT97_28395 | |
| KYT97_28775 | |
| K6K13_01025 | *S. purcellii* |
| K6K13_01055 | |
| K6K13_14675 | |
| K6K13_14710 | |
| K6K13_14780 | |
| K6K13_15505 | |
| K6K13_15750 | |
| K6K13_15755 | |
| K6K13_15760 | |
| K6K13_17015 | |

| | | |
|---|---|---|
| K6K13_17860 | | |
| K6K13_17985 | | |
| K6K13_17990 | | |
| K6K13_18065 | | |
| K6K13_18070 | | |
| K6K13_18395 | | |
| K6K13_18495 | | |
| K6K13_20225 | | |
| K6K13_20255 | | |
| K6K13_20595 | | |
| K6K13_22305 | | |
| K6K13_22335 | | |
| K6K13_22340 | | |
| K6K13_22345 | | |
| K6K13_22350 | | |
| K6K13_22595 | | |
| LY180_00050 | *E. coli* | Folate |
| LY180_00245 | | |
| LY180_00690 | | |
| LY180_02565 | | |
| LY180_04110 | | |
| LY180_04115 | | |
| LY180_04120 | | |
| LY180_04130 | | |
| LY180_04360 | | |
| LY180_05685 | | |

| | |
|---|---|
| LY180_08375 | |
| LY180_09435 | |
| LY180_09710 | |
| LY180_11255 | |
| LY180_11935 | |
| LY180_11995 | |
| LY180_14050 | |
| LY180_14105 | |
| LY180_14200 | |
| LY180_14805 | |
| LY180_15785 | |
| LY180_16410 | |
| LY180_17235 | |
| LY180_19975 | |
| ArsFIN_03620 | *A. nasoniae* |
| ArsFIN_05450 | |
| ArsFIN_05640 | |
| ArsFIN_06090 | |
| ArsFIN_15630 | |
| ArsFIN_15640 | |
| ArsFIN_15660 | |
| ArsFIN_15890 | |
| ArsFIN_15940 | |
| ArsFIN_16650 | |
| ArsFIN_17520 | |
| ArsFIN_24930 | |

| | |
|---|---|
| ArsFIN_26710 | |
| ArsFIN_34690 | |
| ArsFIN_35460 | |
| ArsFIN_36710 | |
| ArsFIN_37470 | |
| ArsFIN_37480 | |
| ArsFIN_41870 | |
| ArsFIN_42040 | |
| ArsFIN_50850 | |
| KYT97_03260 | *R. globerulus* |
| KYT97_07165 | |
| KYT97_07170 | |
| KYT97_07175 | |
| KYT97_07180 | |
| KYT97_09325 | |
| KYT97_14690 | |
| KYT97_16325 | |
| KYT97_19645 | |
| KYT97_22510 | |
| KYT97_22515 | |
| KYT97_22520 | |
| KYT97_22525 | |
| KYT97_25005 | |
| KYT97_26120 | |
| KYT97_27965 | |
| KYT97_28125 | |

| | |
|---|---|
| KYT97_29145 | |
| KYT97_29475 | |
| K6K13_03270 | *S. purcellii* |
| K6K13_05595 | |
| K6K13_07660 | |
| K6K13_07750 | |
| K6K13_07755 | |
| K6K13_07760 | |
| K6K13_07770 | |
| K6K13_08515 | |
| K6K13_10640 | |
| K6K13_11570 | |
| K6K13_12770 | |
| K6K13_14140 | |
| K6K13_14690 | |
| K6K13_15745 | |
| K6K13_16485 | |
| K6K13_16985 | |
| K6K13_16990 | |
| K6K13_17140 | |
| K6K13_18170 | |
| K6K13_18280 | |
| K6K13_18595 | |
| K6K13_19140 | |
| K6K13_21090 | |
| K6K13_21490 | |

| | | |
|---|---|---|
| LY180_21085 | *E. coli* | Cobalamin/ Methionine |
| LY180_15010 | *E. coli* | Cobalamin |
| LY180_15015 | | |
| KYT97_03700 | *R. globerulus* | |
| KYT97_03705 | | |
| KYT97_03710 | | |
| KYT97_04635 | | |
| KYT97_14220 | | |
| KYT97_26700 | | |
| LY180_00550 | *E. coli* | Lipoic Acid |
| LY180_00555 | | |
| LY180_00560 | | |
| LY180_03395 | | |
| LY180_03405 | | |
| LY180_03830 | | |
| LY180_03835 | | |
| LY180_14945 | | |
| LY180_14950 | | |
| LY180_14955 | | |
| LY180_23020 | | |
| ArsFIN_10280 | *A. nasoniae* | |
| ArsFIN_10290 | | |
| ArsFIN_12220 | | |
| ArsFIN_12230 | | |

| | |
|---|---|
| ArsFIN_16560 | |
| ArsFIN_31700 | |
| ArsFIN_31900 | |
| ArsFIN_31910 | |
| ArsFIN_31920 | |
| KYT97_00905 | |
| KYT97_04810 | |
| KYT97_04835 | |
| KYT97_06760 | |
| KYT97_06775 | |
| KYT97_06785 | |
| KYT97_06790 | |
| KYT97_10890 | *R. globerulus* |
| KYT97_16450 | |
| KYT97_23570 | |
| KYT97_23575 | |
| KYT97_25470 | |
| KYT97_27065 | |
| KYT97_27070 | |
| KYT97_27075 | |
| KYT97_27780 | |
| K6K13_03515 | *S. purcellii* |
| K6K13_03520 | |
| K6K13_03525 | |
| K6K13_06115 | |
| K6K13_06120 | |

| | | |
|---|---|---|
| K6K13_06430 | | |
| K6K13_06435 | | |
| K6K13_12630 | | |
| K6K13_17795 | | |
| K6K13_17800 | | |
| K6K13_17805 | | |
| WCLE_002580 | *W. C. lectularius* | |
| WCLE_002730 | | |
| WCLE_003200 | | |
| WCLE_004220 | | |
| WCLE_007840 | | |
| WCLE_00840 | | |
| WCLE_009650 | | |
| WCLE_010890 | | |
| WCLE_011520 | | |
| LY180_00165 | *E. coli* | Arginine |
| LY180_00170 | | |
| LY180_01990 | | |
| LY180_02765 | | |
| LY180_02935 | | |
| LY180_04875 | | |
| LY180_09170 | | |
| LY180_11865 | | |
| LY180_14470 | | |
| LY180_14790 | | |
| LY180_16385 | | |

| | |
|---|---|
| LY180_17230 | |
| LY180_20055 | |
| LY180_20765 | |
| LY180_20770 | |
| LY180_20775 | |
| LY180_20780 | |
| LY180_22335 | |
| KYT97_01240 | *R. globerulus* |
| KYT97_01725 | |
| KYT97_02335 | |
| KYT97_02340 | |
| KYT97_03135 | |
| KYT97_03140 | |
| KYT97_05100 | |
| KYT97_05105 | |
| KYT97_05115 | |
| KYT97_05120 | |
| KYT97_05125 | |
| KYT97_05130 | |
| KYT97_05135 | |
| KYT97_06805 | |
| KYT97_11140 | |
| KYT97_11625 | |
| KYT97_11900 | |
| KYT97_15605 | |
| KYT97_17295 | |

| | |
|---|---|
| KYT97_19975 | |
| KYT97_20915 | |
| KYT97_20920 | |
| KYT97_20925 | |
| KYT97_21010 | |
| KYT97_24860 | |
| KYT97_25090 | |
| KYT97_25095 | |
| KYT97_25100 | |
| KYT97_25335 | |
| KYT97_26290 | |
| KYT97_26400 | |
| KYT97_26410 | |
| KYT97_29080 | |
| K6K13_00720 | *S. purcellii* |
| K6K13_01675 | |
| K6K13_04700 | |
| K6K13_04705 | |
| K6K13_04710 | |
| K6K13_04715 | |
| K6K13_07100 | |
| K6K13_11830 | |
| K6K13_14020 | |
| K6K13_15010 | |
| K6K13_18195 | |
| K6K13_18200 | |

| | | |
|---|---|---|
| K6K13_19270 | | |
| K6K13_20450 | | |
| K6K13_20740 | | |
| K6K13_20990 | | |
| K6K13_21485 | | |
| LY180_00010 | | Methionine |
| LY180_08460 | | |
| LY180_15560 | | |
| LY180_17620 | | |
| LY180_19845 | *E. coli* | |
| LY180_20670 | | |
| LY180_20675 | | |
| LY180_21055 | | |
| LY180_21125 | | |
| Pd630_LPD00737 | *R. opacus* | |
| Pd630_LPD00738 | | |
| Pd630_LPD01440 | | |
| Pd630_LPD02472 | | |
| Pd630_LPD02887 | | |
| Pd630_LPD04089 | | |
| Pd630_LPD04943 | | |
| Pd630_LPD05 | | |

| | |
|---|---|
| 618 | |
| Pd630_LPD06299 | |
| K6K13_00110 | |
| K6K13_00840 | |
| K6K13_00845 | |
| K6K13_01600 | |
| K6K13_01690 | |
| K6K13_04675 | *S. purcellii* |
| K6K13_04850 | |
| K6K13_04890 | |
| K6K13_04935 | |
| K6K13_18310 | |
| K6K13_21970 | |
